# nf-sarcopipe enables integrative discovery of exercise-responsive miRNAs and miRNA–mRNA regulatory networks associated with skeletal muscle adaptation

**DOI:** 10.64898/2026.08.12.744488

**Authors:** Natalia Poblete-Durán, Felipe Gómez-Molina, Gabriel Cabas-Mora, Alex Di Genova-Bravo, Denisse Valladares-Ide, Carol Moraga-Quinteros

**Author notes:** Corresponding authors: Dr. Carol Moraga-Quinteros, Dr. Denisse Valladares-Ide.

## Abstract

Skeletal muscle dynamically adapts to physiological stimuli such as exercise through coordinated molecular and structural remodeling processes. Circulating microRNAs (miRNAs) represent promising non-invasive biomarkers of exercise responsiveness and skeletal muscle physiological states; however, most analytical frameworks rely solely on annotated miRNAs and overlook novel candidates. Here, we present nf-sarcopipe, a modular Nextflow pipeline that integrates *de novo* and reference-guided miRNA discovery with transcriptomic analysis and regulatory network reconstruction. The pipeline is organized into three complementary modules: 1) Preprocessing, 2) miRNA Discovery, and 3) Target Prediction & mRNA Integration. Using publicly available datasets from active and sedentary young women, the pipeline identified reproducible miRNA signatures and prioritized a small set of structurally supported, high-confidence *de novo* candidates. Previously reported exercise-associated miRNAs compiled from the literature were additionally incorporated for comparative candidate evaluation. Although the available datasets were derived from different tissues, confounding-aware analyses enabled the identification of coherent transcriptional signatures associated with exercise responsiveness. Integrative miRNA–mRNA analysis uncovered consistent regulatory interactions linking circulating miRNAs—both novel and known—to pathways involved in immune response, extracellular matrix remodeling, autophagy, and skeletal muscle adaptation. Together, these results establish nf-sarcopipe as a robust and scalable framework for complementary miRNA discovery and for investigating regulatory mechanisms associated with exercise-induced skeletal muscle adaptation.

## Introduction

Skeletal muscle is a highly adaptable tissue that dynamically responds to physiological stimuli such as exercise ^1^. This adaptive capacity is essential for maintaining muscle function, metabolic homeostasis, mobility, and overall systemic health ^1,2^. Regular exercise promotes coordinated molecular and structural remodeling processes, including mitochondrial biogenesis, angiogenesis, protein turnover, and neuromuscular adaptation, which collectively support muscle performance and resilience ^3,4^. In contrast, sedentary behavior has been associated with impaired muscle function, metabolic dysregulation, and altered intracellular signaling pathways, contributing to detrimental changes in skeletal muscle physiology ^4,5^. Characterizing the molecular mechanisms underlying these adaptive and maladaptive responses is therefore critical to identify early regulatory changes and develop strategies to preserve muscle health across different physiological conditions ^6^. Because many of these adaptive pathways are also disrupted during muscle aging and disease, understanding post-transcriptional regulation may facilitate the identification of biomarkers and therapeutic targets relevant to skeletal muscle disorders, including sarcopenia ^7^.

Among the molecular regulators involved in skeletal muscle adaptation, microRNAs (miRNAs) have emerged as key post-transcriptional modulators of gene expression ^8^. miRNAs are small non-coding RNAs capable of regulating messenger RNAs (mRNAs) through translational repression or transcript degradation, thereby influencing multiple biological processes including satellite cell activation, myogenesis, angiogenesis, and metabolic adaptation ^9–11^. Early studies identified muscle-enriched miRNAs such as miR-1, miR-133 ^12,13^), and miR-206 ^14^ as critical regulators of skeletal muscle proliferation and differentiation, while miR-146 was initially associated with inflammatory regulatory pathways ^13^. Subsequent studies demonstrated that several of these miRNAs are responsive to exercise in both skeletal muscle tissue and circulation, including miR-1, miR-21, miR-133, miR-146, and miR-206, supporting their role in systemic adaptation and inter-tissue communication ^15–17^. However, exercise-responsive miRNA profiles can vary according to exercise modality, intensity, duration, and chronicity, highlighting the complexity of the underlying regulatory landscape ^17,18^.

Importantly, circulating miRNAs reflect dynamic regulatory processes associated with skeletal muscle adaptation to exercise, including pathways involved in muscle remodeling, regeneration, and metabolic regulation ^15^. Changes in the expression of specific miRNAs have been linked to muscle regeneration, anabolic signaling, inflammation, and metabolic regulation, and may precede measurable alterations in muscle mass or physiological performance ^19,20^. Due to their sensitivity to physiological stimuli, circulating miRNAs have gained increasing attention as minimally invasive biomarkers capable of distinguishing different physiological states, including active and sedentary conditions ^16,21^.

Current computational approaches for small RNA sequencing analysis predominantly rely on reference-guided strategies based on alignment against annotated miRNA databases such as miRBase ^22^. In these approaches, sequencing reads are typically mapped to previously annotated miRNA repertoires, prioritizing the detection and quantification of known miRNAs while potentially excluding unannotated or condition-specific candidates. As a result, the discovery of novel miRNAs associated with particular physiological conditions may remain limited, especially in dynamic processes such as exercise-induced skeletal muscle remodeling ^23,24^. Although several studies have characterized exercise-responsive miRNAs using conventional annotation-dependent workflows, the regulatory landscape underlying skeletal muscle adaptation remains incompletely understood, particularly considering the potential contribution of dynamic and condition-specific regulatory molecules to inter-individual variability and adaptive responses ^25^. In addition, computational target prediction frequently produces large numbers of candidate miRNA–mRNA interactions, many of which represent false positives due to reliance on sequence complementarity alone. Consequently, integrating sequence-based prediction with coordinated expression patterns provides a robust strategy for prioritizing biologically meaningful miRNA–mRNA interactions and reconstructing post-transcriptional regulatory networks ^26,27^.

To address these limitations, we developed nf-sarcopipe, a modular pipeline implemented in Nextflow ^28^ designed to improve the discovery of novel miRNAs and reconstruct biologically meaningful post-transcriptional regulatory networks through the integration of small RNA-seq and RNA-seq datasets. The pipeline (Figure 1) is organized into three complementary modules: 1) Preprocessing, 2) miRNA Discovery, and 3) Target Prediction & mRNA Integration. The Preprocessing module performs quality control and complementary *de novo* and reference-guided miRNA detection using BrumiR ^29^ and miRDeep2 ^23^ maximizing the identification of both uncharacterized and annotated miRNAs. The miRNA Discovery module reduces redundant candidate sequences from BrumiR-core output, incorporates structural validation, and generates expression matrices to improve confidence in candidate miRNA annotation and downstream expression analyses. Finally, the Target Prediction & mRNA Integration module performs differential expression analysis, transcriptome-wide target prediction based on seed complementarity, prioritization of coherent miRNA–mRNA interactions through inverse expression patterns, functional enrichment, and integration with mRNA-seq data to reconstruct biologically meaningful post-transcriptional regulatory networks.

**Figure 1.**
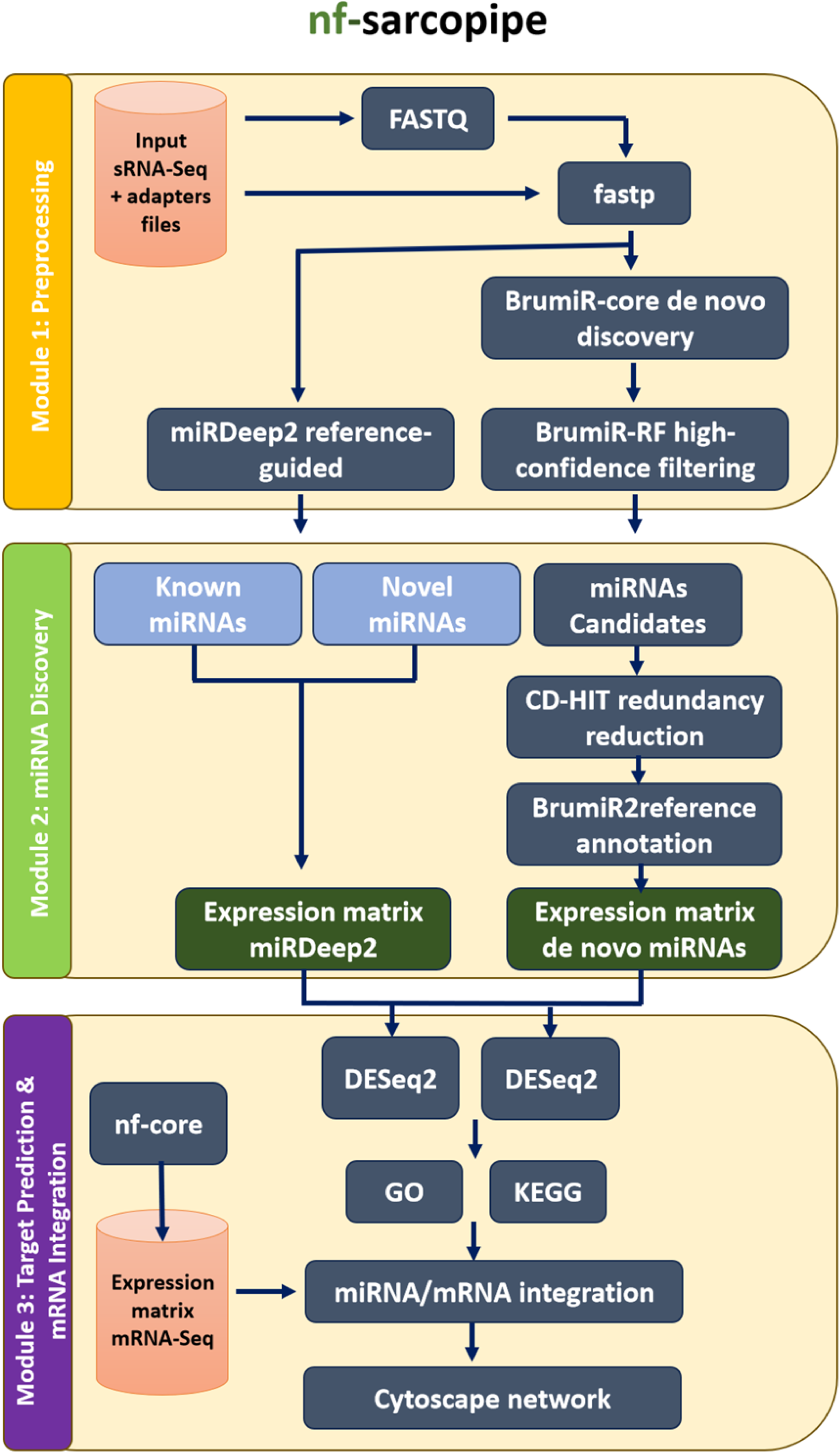
Overview of the nf-Sarcopipe workflow. Sarcopipe is a modular Nextflow DSL2 pipeline designed for the integrated analysis of sRNA-seq and mRNA-seq data to identify miRNA-mRNA regulatory interactions associated with exercise adaptation and skeletal muscle remodeling. The workflow is organized into three modules: (I) Preprocessing, which includes quality control and adapter trimming ***(fastp), de novo*** miRNA discovery using BrumiR, machine learning-based filtering (BrumiR-RF), and reference-guided detection using miRDeep2; (II) miRNA Discovery, which performs redundancy reduction (CD-HIT), genomic annotation (BrumiR2Reference), expression matrix construction, and identification of core miRNAs based on intra-group presence; and (III) Target Prediction & mRNA Integration, which includes differential expression analysis (DESeq2) and integrative miRNA-mRNA network reconstruction using mRNA-seq expression data.

Using publicly available datasets from active and sedentary young women as a pilot framework, including circulating small RNA-seq data ^30^, and complementary RNA-seq datasets from age-matched active and sedentary cohorts ^31,32^, we demonstrate how this integrative strategy enables the systematic exploration of regulatory networks associated with exercise responsiveness. To facilitate biological interpretation, previously reported exercise-associated miRNAs were systematically curated from the literature and incorporated as a reference resource for comparison with candidate miRNAs identified by the pipeline (Supplementary Tables 7; page 1 & 2). Given the heterogeneous tissue origins of the transcriptomic datasets, confounding-aware analyses were incorporated to prioritize exercise-associated regulatory signatures. This approach provides a reproducible framework for the identification and characterization of both novel and annotated miRNAs and their associated regulatory networks in skeletal muscle. Although demonstrated here using exercise-responsive datasets from healthy young individuals, the pipeline is readily applicable to future studies investigating skeletal muscle remodeling under physiological and pathological conditions, including aging, muscle wasting, and sarcopenia, where integrated miRNA–mRNA analyses may facilitate the discovery of regulatory mechanisms and candidate biomarkers.

## Results

### 1. -Robust miRNA candidate discovery from public plasma sRNA-seq data in active and sedentary young women

nf-sarcopipe was designed to process raw sRNA-seq data through sequential steps of quality control, size selection, and parallel miRNA discovery (Figure 1). Here the pipeline integrates both *de novo* (BrumiR) and reference-guided (miRDeep2) strategies, allowing the same dataset to be interrogated through complementary miRNAs discovery approaches.

We collected a publicly available plasma sRNA-seq dataset from active and sedentary young women (see Materials and Methods, Section 1) to demonstrate the utility and capabilities of Module I of nf-sarcopipe (Figure 2a). The dataset includes 25 samples (10 active synchronized swimmers prior to exercise and 15 sedentary controls), that among other uses provides a suitable framework to evaluate preprocessing performance. Across all samples, sequencing depth ranged from 11.7 to 29.3 million reads per sample, totaling 530.2 million reads (Supplementary Table 1). Adapter trimming and quality filtering with fastp retained 226.4 million reads (43.8% of the raw input). The subsequent size-selection step (18–28 nt interval defined for miRNA discovery) reduced this to 208.4 million reads (37.8% of total reads), consistent with enrichment for short RNA species. FastQC and *fastp* quality reports indicated that read loss was primarily attributable to adapter-trimmed reads falling below the minimum length threshold, whereas fewer than 0.002% of reads were discarded because of low sequencing quality (Supplementary Table 2).

**Figure 2.**
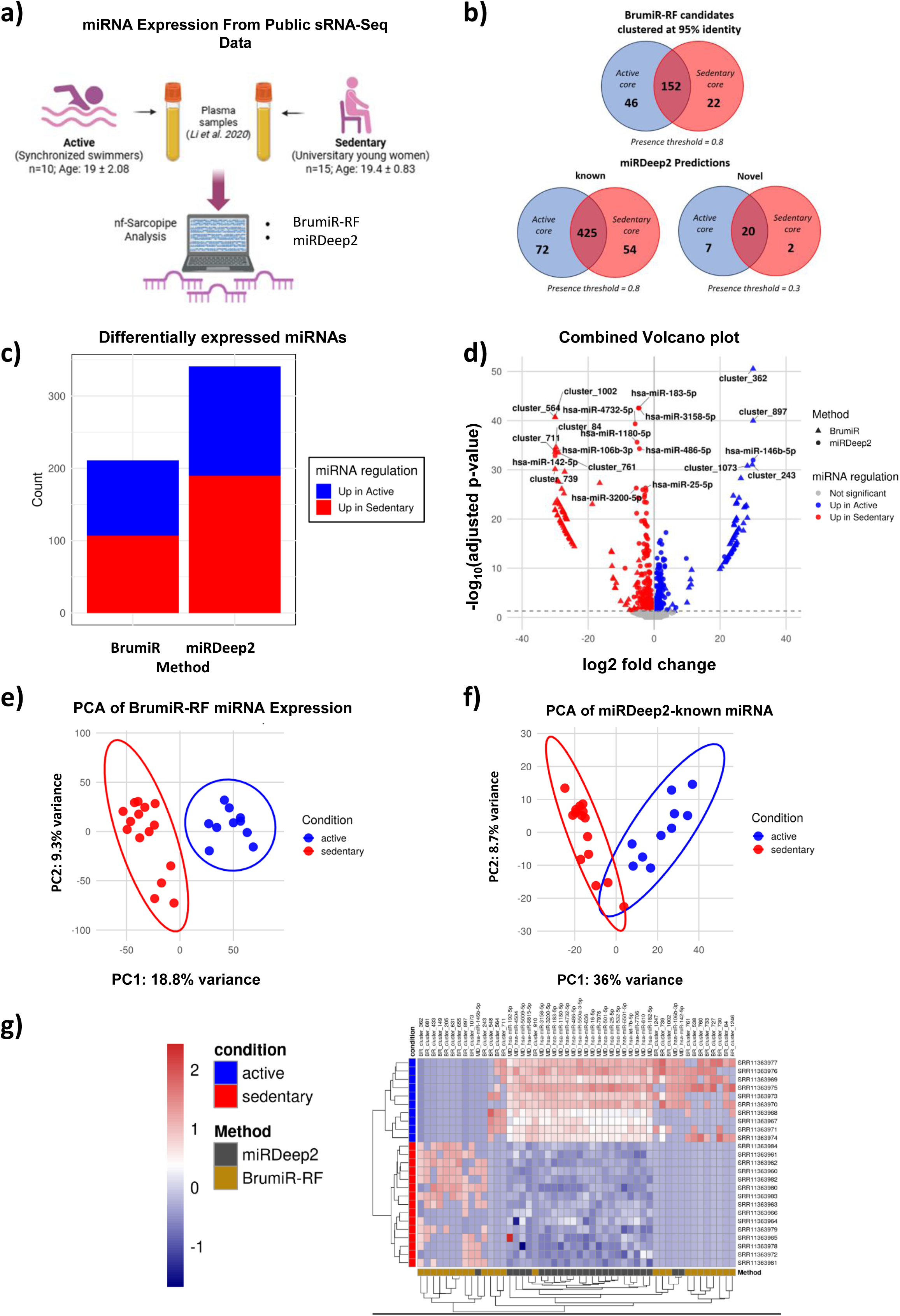
Overview and statistical characterization of circulating miRNA expression profiles analyzed with nf-Sarcopipe. (a) Schematic representation of the study design and data processing workflow. Public plasma sRNA-seq data from active (synchronized swimmers, n = 10) and sedentary (university women, n = 15) individuals were processed using nf-Sarcopipe, integrating both reference-guided (miRDeep2) and ***de novo*** (BrumiR-RF) miRNA detection strategies, (b) Venn diagrams summarizing core miRNA sets for each method and condition. For BrumiR-RF, candidate miRNAs clustered at 95% sequence identity were filtered using a presence threshold of 0.8. For miRDeep2, known miRNAs (miRBase-annotated) were filtered at a presence threshold of 0.8, while novel predictions were filtered at 0.3. Presence thresholds were selected to ensure sufficient representation of detectable miRNA features across samples while maintaining consistency between methods; accordingly, BrumiR-RF candidates were analyzed using a presence threshold of 0.8. miRDeep2 novel predictions were not considered in downstream analyses, (c) Stacked bar plot showing the number of differentially expressed miRNAs identified by each method. Differential expression analysis was performed using DESeq2 with Wald test statistics and Benjamini-Hochberg correction for multiple testing (adjusted p < 0.05). (d) Combined volcano plot displaying Iog2 fold change versus -Iog10 adjusted p-value for miRNAs detected by BrumiR-RF and miRDeep2. Ibp 10 Significant miRNAs from different algorithms are colored according to regulation direction, and point shapes indicate the detection method, (e-f) Principal component analysis (PCA) of variance-stabilized miRNA expression values (DESeq2 VST transformation). PCA was performed independently for (e) BrumiR-RF candidates and miRDeep2-known miRNAs (f) using the top 500 most variable features, (g) Heatmap hierarchical clustering of the 25 most significant BrumiR-RF candidate clusters and the 25 most significant miRDeep2 annotated miRNAs, using variance-stabilized expression values scaled by feature.

BrumiR-core processed a total of 208.4 million reads within the canonical miRNA size range (18–28 nt) and 17.98 million reads exceeding 28 nt (Supplementary Table 1). Analysis of the 18–28 nt fraction resulted in the assembly of 1,624,439 unitigs, 18,280 additional sequences, and 12,987 raw candidate miRNAs, of which 8,713 were retained as high-confidence candidates following BrumiR-RF filtering in where 3,346 are unique sequences (Supplementary Table 1). In parallel, miRDeep2 generated 19,118 annotated miRNA detection records corresponding to 3,461 unique mature annotated sequences, together with 30,402 novel prediction records representing 5,868 unique mature candidate sequences across all sequencing libraries (Supplementary Table 1). These values do not represent unique human miRNA genes; rather, they correspond to cumulative detections across samples, including repeated observations of annotated mature miRNAs. Similarly, novel miRNA predictions are made independently for each sequencing library, so the same mature sequence may be detected in multiple samples or predicted from different precursor sequences. As miRDeep2 is designed to maximize sensitivity during novel miRNA discovery, it intentionally generates a broad initial catalogue of candidates that is expected to be subsequently refined through additional biological and computational prioritization steps ^33^. For example, the current human miRBase release (v22.1) contains 1,917 annotated precursor miRNAs that produce 2,654 mature miRNA sequences ^34^. At the sample level, BrumiR-RF (Random Forest) consistently recovered hundreds of candidates per sample, whereas miRDeep2 identified on the order of one thousand annotated miRNA detection records per library. These differences reflect the distinct strategies implemented by each algorithm: BrumiR performs reference-free *de novo* reconstruction of small RNA candidates, whereas miRDeep2 leverages genome alignment and annotated precursor hairpin structures to identify known miRNAs while also predicting novel candidates. Accordingly, the large number of initial novel predictions represents the expected high-sensitivity output of miRDeep2 rather than a final set of bona fide novel miRNAs. In the present pipeline, these candidates are subsequently subjected to clustering, cross-sample reproducibility filtering, differential expression analysis, structural prioritization, and functional integration with mRNA-seq data to obtain a small set of high-confidence novel miRNA candidates. Consequently, the two approaches provide complementary views of the miRNA landscape rather than redundant predictions. Overall, these results show that Module I effectively processes raw plasma sRNA-seq data and generates a diverse set of miRNA candidates, establishing the basis for downstream analyses.

### 2. -Reproducible miRNA core sets identification through integrated *de novo* and reference-guided strategies

To identify reproducible miRNAs suitable for downstream analyses, nf-sarcopipe retained only candidates consistently detected across individuals. Throughout this study, reproducibility was defined as detection in at least 80% of the samples within each study group (presence ≥0.8). For *de novo* miRNAs, clustering analyses showed that increasing sequence identity stringency or requiring detection in all samples reduced the total number of retained candidates but did not reveal additional reproducible miRNAs (Supplementary Figure 1a,b). Therefore, a clustering threshold of 95% sequence identity together with the default presence criterion (≥0.8) was selected, providing the most confident dataset while preserving the complete set of reproducible candidates (Figure 2b).

The same reproducibility criterion was applied to the reference-guided branch. Under this threshold, annotated miRNAs were consistently detected across samples, whereas most novel miRDeep2 predictions showed limited reproducibility. Novel candidates were recovered only after relaxing the presence threshold to 0.3, indicating that they were detected in only a minority of individuals (Figure 2b; Supplementary Figure 1c). We decided to continue exclusively on annotated (miRBase) miRNAs of miRDeep2 to prioritize robust and reproducible miRNA candidates.

Differential expression analysis identified slightly more miRNAs upregulated in sedentary than in active individuals with both discovery strategies (Figure 2c). BrumiR-RF detected 211 differentially expressed candidate clusters (107 sedentary, 104 active), whereas miRDeep2 identified 341 annotated miRNAs (190 sedentary, 151 active). Both methods showed comparable effect size and significance distributions in the combined volcano plot (Figure 2d) (see Supplementary Figures S1D and S1E for volcano plots generated separately for each algorithm). The threshold selected to principal component analysis (PCA) was using the 500 most variable miRNAs because the filtered expression matrices contained 1,307 BrumiR-RF candidates and 861 miRDeep2 known candidates, allowing the same feature-selection criterion to be applied to both discovery strategies while focusing on the most informative expression patterns. Samples were separated into active and sedentary groups for both BrumiR-RF (PC1 = 18.8%, PC2 = 9.3%) and miRDeep2 (PC1 = 36.0%, PC2 = 8.7%) (Figure 2e–f).

Hierarchical clustering of the 25 most significant features from each method also grouped samples according to condition (Figure 2g), supporting the reproducibility of the identified miRNA expression patterns.

### 3. - Structural and seed-based prioritization identifies a high-confidence set of *de novo* miRNAs

In the following step, nf-Sarcopipe prioritized reproducible BrumiR-RF-derived candidates through sequential structural validation, reference annotation, and seed-based characterization (Figure 3a). Structural validation with BrumiR2Reference reduced the initial BrumiR-RF core set from 46 active-specific, 22 sedentary-specific, and 152 shared clusters to 5, 2, and 20 structurally supported candidates, respectively (Figure 3b, Supplementary Table 3), highlighting the importance of secondary-structure filtering for reducing potential false-positive predictions.

**Figure 3.**
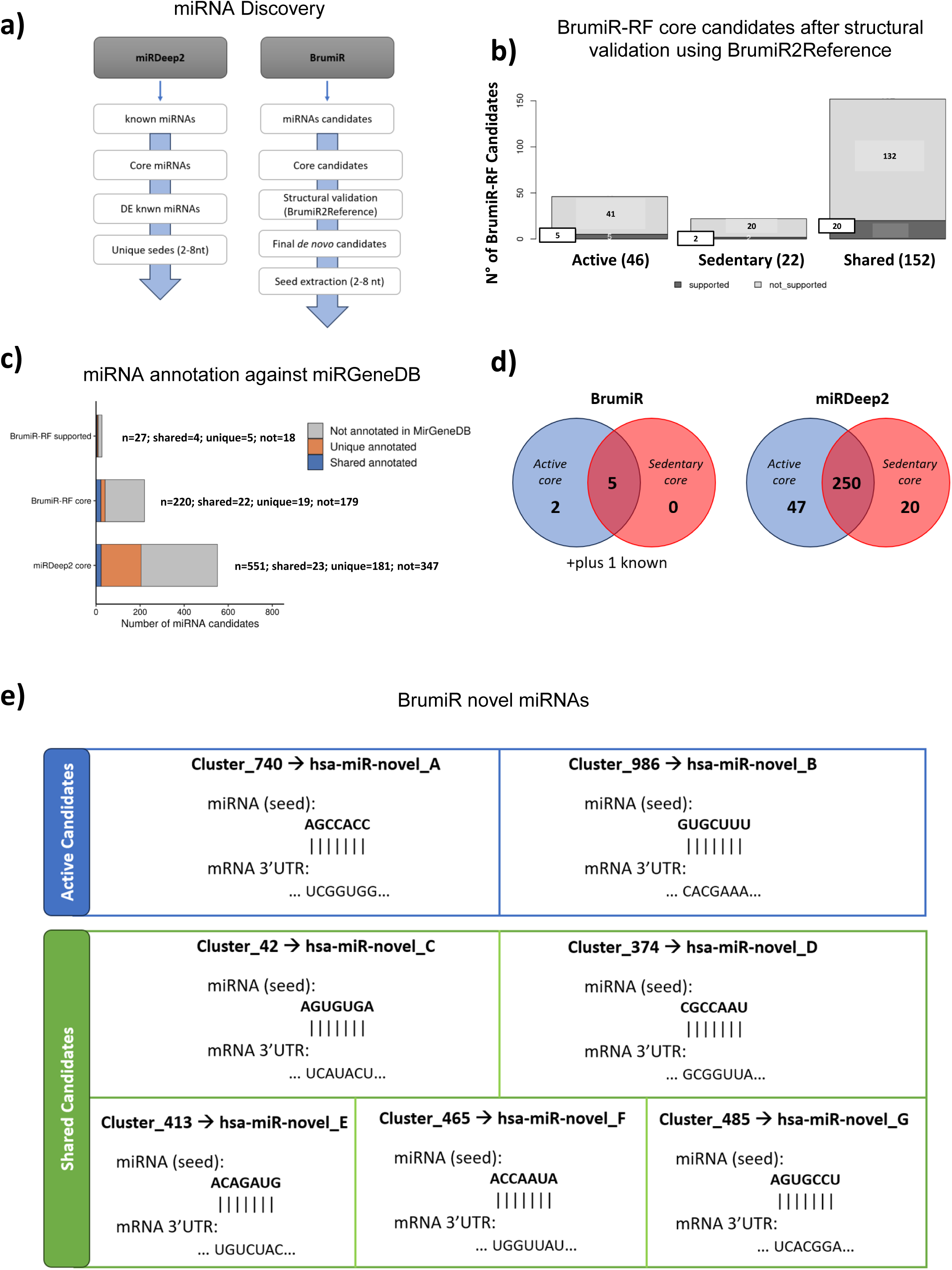
Prioritization and functional characterization of de novo miRNA candidates identified by nf-Sarcopipe. (a) Overview of the miRNA discovery and prioritization workflow. miRDeep2-derived miRNAs were subjected to differential expression analysis followed by extraction of seed regions (nucleotides 2-8), yielding a set of unique seeds for downstream target prediction. In parallel, BrumiR-derived candidates were filtered through core set definition, structural validation (BrumiR2Reference), and seed-based target prediction, resulting in a refined set of high-confidence ***de novo*** miRNA candidates, (b) Stacked bar plot summarizing BrumiR-RF core candidates (presence ≥0.8) across conditions (active, sedentary, and shared) following structural validation using BrumiR2Reference. Dark and light segments indicate candidates supported or not supported by genomic mapping and secondary structure prediction (RNAfold), respectively, (c) Annotation of BrumiR-RF and miRDeep2 core miRNA candidates against MirGeneDB. Matches required ≥98% identity, ≥75% query coverage, and zero mismatches. BrumiR-RF candidates are shown before and after BrumiR2Reference structural validation. miRDeep2 candidates classified as ***Not annotated in MirGeneDB*** correspond to known miRNAs annotated in miRBase but lacking a qualifying MirGeneDB match, (d) Distribution of final differentially expressed miRNA candidates according to their original core-set membership. BrumiR-RF candidates correspond to structurally supported ***de novo*** miRNAs with adjusted p < 0.05, whereas miRDeep2 candidates correspond to differentially expressed annotated miRNAs within the known core union. Shared candidates were recurrently detected in both groups but differed significantly in abundance, (e) Schematic representation of seed-mediated target interactions for representative BrumiR-derived miRNA ***de novo*** candidates. Seed regions (nucleotides 2-8) are aligned to complementary sequences within predicted mRNA 3’UTRs, illustrating canonical base-pairing interactions. Candidate clusters were renamed using a standardized nomenclature (hsa-miR-novel_A-G).

Annotation against MirGeneDB further classified the retained candidates. Among the 220 BrumiR-RF core candidates, 22 corresponded to previously annotated miRNAs, whereas 179 remained unannotated. Within the structurally supported subset, four candidates matched annotated miRNAs and five represented unique annotated sequences, while 18 lacked annotation and were therefore retained as high-confidence *de novo* candidates. In contrast, the miRDeep2 core set contained 551 annotated miRNAs, including 23 shared with BrumiR-RF and 181 uniquely detected by the reference-guided approach, whereas 347 candidates lacked MirGeneDB annotation (Figure 3c). Because all annotated BrumiR-RF candidates were represented by their corresponding miRDeep2 annotations, only structurally supported *de novo* candidates were retained from the BrumiR branch for downstream analyses.

Following differential expression analysis, seven *de novo* miRNAs were retained from the final BrumiR candidate set, including two from the active-specific core and five from the shared core, together with one active-specific known miRNA (hsa-miR-660-5p) that was also identified by the miRDeep2 annotation (Figure 3d). In parallel, the final miRDeep2 dataset comprised 317 differentially expressed annotated miRNAs, including 47 from the active-specific core, 250 from the shared core, and 20 from the sedentary-specific core (Figure 3d).

The seven final *de novo* miRNAs candidates originated from clusters containing 9–32 highly similar sequences, some of which may represent putative isomiRs (Supplementary Figure 2a–b). Predicted precursor secondary structures were consistent with canonical miRNA hairpins and provided independent structural support for all retained candidates (Supplementary Figure 2c). Canonical seed inspection enabled standardized nomenclature (hsa-miR-novel_A to hsa-miR-novel_G) and confirmed perfect seed complementarity with predicted target sites (Figure 3e). Comprehensive 7-mer seed-space analysis further demonstrated that only two candidates (hsa-miR-novel_E and hsa-miR-novel_G) shared canonical seed sequences with previously annotated human miRNA families. The remaining five candidates contained novel canonical seed sequences but showed isolated non-canonical 7-mer similarities elsewhere within the mature sequence (Supplementary Figure 2d; Supplementary Table 4). Notably, hsa-miR-novel_A showed no canonical or non-canonical similarity to any known human miRNA seed, further supporting its potential novelty.

### 4. - Correction of tissue-driven confounding reveals a high-confidence transcriptional signature and supports integrative miRNA–mRNA regulatory analysis

Following confounding correction using a principal component-based linear model from whole blood in active and vastus lateralis biopsies (Figure 4a; see Methods: mRNA-seq processing, confounding correction and integrative analysis), differential expression analysis identified a high-confidence transcriptional signature comprising 36 genes (Figure 4b–c). PCA of the corrected dataset confirmed separation that was consistent according to biological condition rather than tissue origin (Supplementary Figure 3a–d).

**Figure 4.**
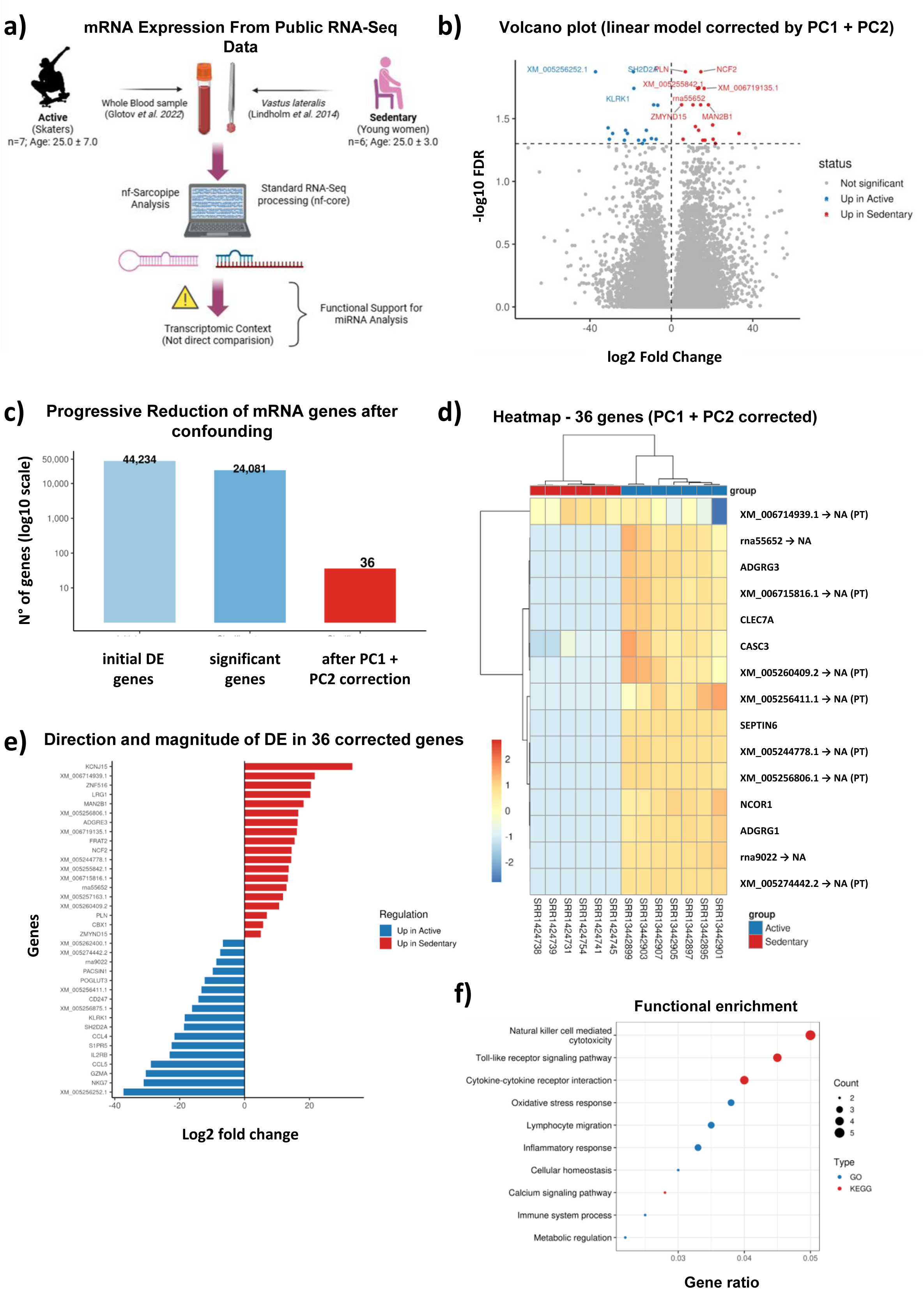
Correction of tissue-driven confounding reveals a robust transcriptional signature associated with physical activity. (a) Schematic overview of the study design, integrating RNA-seq data from whole blood (Active group) and skeletal muscle (Sedentary group), processed through a standardized pipeline, (b) Volcano plot of differential expression results after correction using a linear model including PC1 and PC2 as covariates. A limited number of genes remain significant (FDR < 0.05), indicating removal of confounding-driven signals, (c) Barplot illustrates the progressive reduction in candidate genes from 44,234 filtered transcripts, to 24,081 significant genes prior to correction, and finally to 36 genes after correction for confounding effects, (d) Heatmap of the corrected gene set (n = 36), showing clear separation between Active and Sedentary individuals. Due to filtering and annotation constraints, a subset of genes (n = 15) is displayed. NA = Not Annotated; PT = Predicted Transcript, (e) Diverging barplot showing the direction and magnitude of differential expression for the 36 corrected genes. Positive values indicate genes upregulated in Sedentary individuals, while negative values indicate genes upregulated in Active individuals, (f) Curated functional enrichment analysis of the 36 corrected genes, highlighting pathways related to immune regulation, oxidative stress, inflammation, and cellular homeostasis. Only biologically relevant terms were retained after manual curation to exclude unrelated disease-or infection-driven pathways.

Analysis of expression directionality revealed distinct biological trends between conditions (Figure 4e). Genes upregulated in active individuals (n = 17) were predominantly associated with immune activation and cytotoxic functions, including KLRK1, NKG7, GZMA, CD247, IL2RB, CCL4, and CCL5, consistent with the enhanced immune surveillance and cytotoxic activity previously associated with regular physical activity ^35,36^. Conversely, genes upregulated in sedentary individuals (n = 19) were mainly related to metabolic regulation, oxidative stress, and cellular homeostasis, including PLN, KCNJ15, NCF2, LRG1, and ZNF516, supporting previous observations linking physical inactivity with metabolic dysregulation and oxidative stress ^37^; ^38,39^. Hierarchical clustering further separated active and sedentary samples, supporting the robustness of the corrected transcriptional signature (Figure 4d).

Functional enrichment analysis highlighted pathways related to immune regulation, cytokine signaling, oxidative stress, calcium signaling, inflammation, and cellular homeostasis after manual removal of disease-associated terms (Figure 4f). Immune deconvolution further indicated that the transcriptional profiles were primarily composed of neutrophils, followed by monocytes and T-cell populations, consistent with the expected composition of peripheral blood.

To enable downstream miRNA–mRNA integration, the corrected gene set was filtered according to transcript robust annotation, retaining 27 genes with annotated 3′UTRs suitable for target prediction. These genes constituted the high-confidence transcriptomic input for the integrative regulatory analyses presented in the following section.

### 5. -nf-sarcopipe integrates miRNA–mRNA profiles to identify coherent regulatory interactions and biologically structured networks

To identify post-transcriptional regulatory interactions associated with physical activity, differentially expressed miRNAs identified by both discovery strategies were integrated with the corrected mRNA expression signature. As illustrated in Figure 5a, coherent regulatory interactions were defined by inverse miRNA–mRNA expression relationships, whereby upregulated miRNAs were associated with downregulated target transcripts and vice versa. Canonical 7-mer seed matching was then used to predict candidate miRNA–mRNA interactions, followed by validation using miRanda for *de novo* candidates and multiMiR for annotated miRNAs, as described in the Materials and Methods (section 4f).

**Figure 5.**
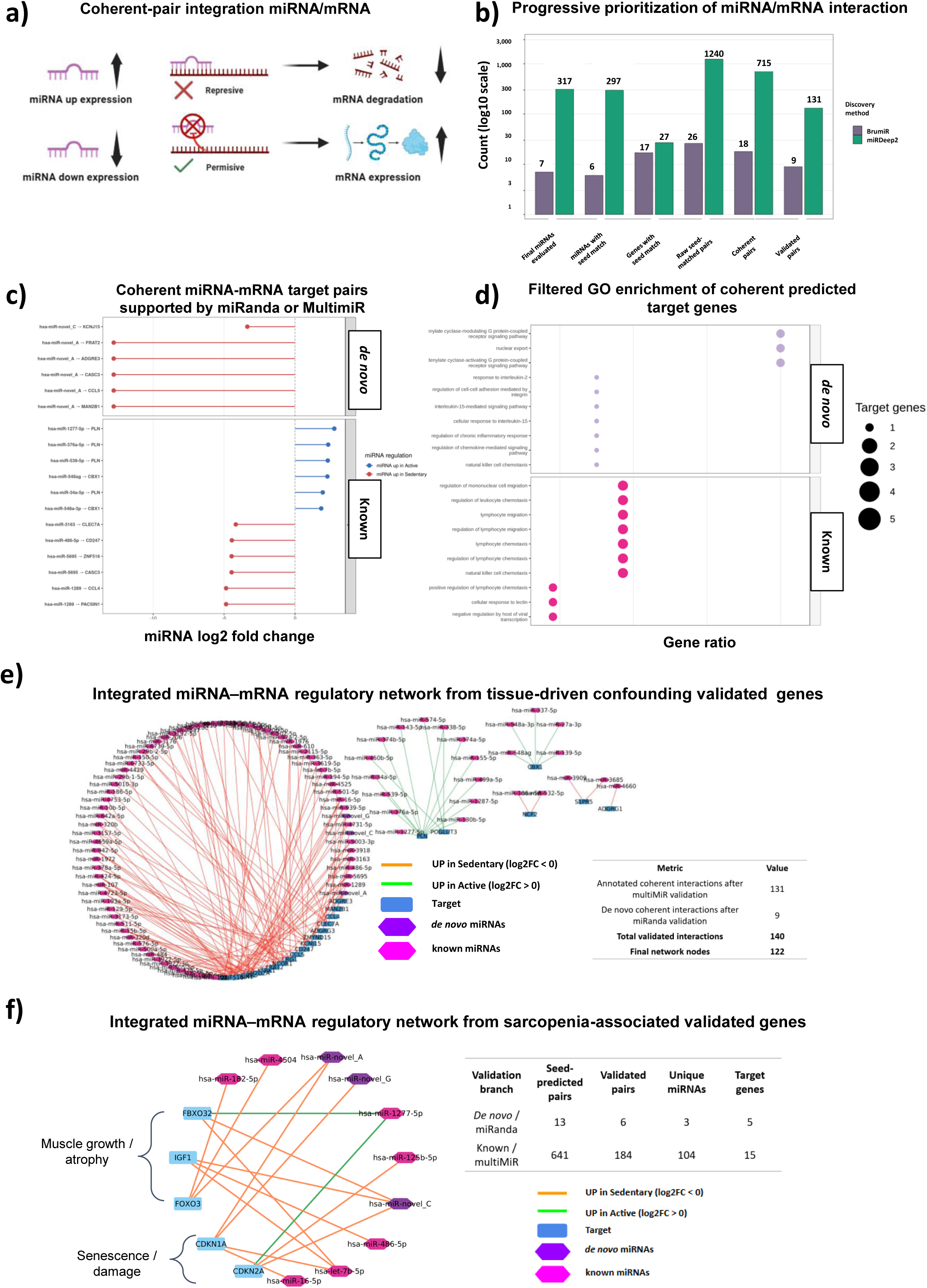
Integrated miRNA-mRNA regulatory landscape combining annotated and *de novo* miRNAs identified by nf-Sarcopipe. (a) Conceptual framework for coherent miRNA-mRNA integration. Inverse expression relationships were used to define biologically coherent interactions, where miRNA upregulation is associated with target mRNA repression and vice versa, (b) Progressive prioritization of miRNA-mRNA interactions identified by nf-Sarcopipe. Sequential reduction of candidate interactions through the target prediction workflow for BrumiR-derived *de novo* miRNAs and miRDeep2 annotated miRNAs. Bars indicate the number of differentially expressed miRNAs evaluated, miRNAs with at least one predicted seed match, target genes containing canonical seed matches, raw seed-matched miRNA-mRNA pairs, coherent interactions exhibiting inverse miRNA-mRNA expression patterns, and independently validated interactions retained after miRanda validation (BrumiR) or multiMiR validation (miRDeep2). The y-axis is displayed on a logarithmic scale, (c) Representative validated coherent miRNA-mRNA interactions. The strongest validated pairs from each discovery branch were ranked according to absolute miRNA Iog2 fold change. BrumiR-derived *de novo* interactions were exclusively associated with miRNAs upregulated in sedentary individuals, whereas validated miRDeep2 interactions included both active-and sedentary-associated miRNAs. (d) Gene Ontology Biological Process enrichment analysis performed separately on validated target genes regulated by de novo and annotated miRNAs. Enriched biological processes were predominantly associated with immune regulation, leukocyte migration, cytokine signaling, and inflammatory responses, reflecting systemic adaptations associated with physical activity, (e) Integrated regulatory network reconstructed from all validated coherent miRNA-mRNA interactions. The network combines interactions supported by miRanda (de *novo* miRNAs) and multiMiR (annotated miRNAs) following seed-based target prediction and inverse-expression filtering. Edge colors indicate whether the corresponding miRNA was upregulated in the active or sedentary group, whereas node colors distinguish annotated miRNAs, *de novo* miRNAs, and target genes. The final high-confidence network comprises 140 validated interactions connecting 122 nodes, (f) Validated interactions between discovered miRNAs and sarcopenia-associated genes. A subset of validated interactions involving genes implicated in muscle growth, atrophy, extracellular matrix remodeling, and cellular senescence is shown. De *novo* miRNAs recovered by BrumiR target canonical regulators including FOXO3, IGF1, FBXO32, CDKN1A, and CDKN2A, illustrating the biological relevance of newly identified candidates within musculoskeletal regulatory pathways.

After differential expression filtering, 317 annotated miRNAs and 7 high-confidence *de novo* miRNA candidates were retained. Seed matching identified predicted interactions for 297 annotated and 6 *de novo* miRNAs, generating 1,240 and 26 putative miRNA–mRNA pairs, respectively. Applying an inverse-expression coherence filter reduced these to 715 coherent pairs involving 27 target genes for annotated miRNAs and 18 coherent pairs involving 13 target genes for *de novo* candidates (Figure 5b).

Subsequent validation with miRanda (*de novo* candidates) and multiMiR (annotated miRNAs) retained 131 annotated and 9 de novo coherent interactions, yielding a final high-confidence regulatory network comprising 140 validated miRNA–mRNA interactions connecting 122 nodes (Figure 5e). This integrated network combines annotated and newly identified miRNAs into a unified regulatory framework associated with physical activity adaptation.

Representative validated coherent miRNA–mRNA interactions are shown in Figure 5c. The strongest validated pairs from each discovery branch were selected according to miRNA log2 fold change. All validated *de novo* interactions were associated with miRNAs upregulated in sedentary individuals, whereas validated known interactions included both active and sedentary-associated miRNAs. Multiple annotated miRNAs converged on common targets, including PLN, CBX1, and CASC3, highlighting shared regulatory hubs within the final network. Notably, PLN is a key regulator of calcium homeostasis and muscle contractility ^40^, whereas CBX1 and CASC3 participate in chromatin organization ^41^ and post-transcriptional RNA regulation ^42^, respectively.

Functional enrichment analysis of the validated target genes identified 22 significantly enriched Gene Ontology Biological Process terms, predominantly associated with immune cell chemotaxis, leukocyte migration, cytokine signaling, and G protein-coupled receptor signaling (Figure 5d). KEGG pathway analysis identified a single significantly enriched pathway, viral protein interaction with cytokine and cytokine receptors, reflecting enrichment of cytokine signaling components including CCL4, CCL5, and IL2RB (Supplementary Figure 4a). Reactome analysis revealed that validated *de novo* miRNA targets were preferentially associated with interleukin-2/interleukin-15 signaling, G protein-coupled receptor signaling, β-catenin-related processes, neutrophil degranulation, and mRNA surveillance, whereas annotated miRNA targets were enriched for chemokine signaling, VEGF signaling, lymphocyte migration, and immune-related pathways (Supplementary Figure 4b). Although none of the Reactome pathways remained significant after multiple-testing correction, these pathway associations support the biological plausibility of the prioritized *de novo* candidates.

To further illustrate the biological utility of the identified miRNAs beyond the dataset used for discovery, we evaluated their predicted regulatory interactions with a curated panel of genes associated with sarcopenia, a progressive musculoskeletal disorder characterized by impaired muscle mass and function. Using the same seed-based prediction strategy followed by validation with miRanda (*de novo* miRNAs) and multiMiR (annotated miRNAs), we identified six validated interactions involving BrumiR-derived *de novo* miRNAs targeting five key musculoskeletal regulators (FOXO3, IGF1, FBXO32, CDKN1A, and CDKN2A), together with 184 validated interactions involving annotated miRNAs and 15 sarcopenia-associated genes (Figure 5f; Supplementary Figure 4c). These results demonstrate that the newly discovered *de novo* miRNAs converge on established regulators of muscle growth, atrophy, and cellular senescence, illustrating how nf-Sarcopipe can generate biologically meaningful candidate miRNAs for downstream studies in musculoskeletal diseases.

## Discussion

Here, we present nf-sarcopipe, a reproducible computational pipeline that integrates *de novo* and reference-guided miRNA discovery with transcriptomic data to reconstruct biologically coherent miRNA–mRNA regulatory networks associated with physical activity. By combining complementary discovery strategies, structural prioritization of *de novo* candidates, differential expression analysis, seed-based target prediction, and complementary validation of predicted interactions using miRanda for *de novo* candidates and multiMiR for annotated miRNAs, the pipeline enables systematic identification of both annotated and previously uncharacterized regulatory miRNAs. The reference-guided branch also recovered numerous well-established exercise-associated miRNAs, including members of the let-7, miR-1, miR-21, miR-29, miR-206, miR-486, and miR-499 families (Supplementary Table 7 page 1 & 2), supporting the biological relevance of the annotated component while complementing it with newly identified *de novo* candidates. In addition, detailed preprocessing reports generated by fastp demonstrated that the substantial reduction in read counts during adapter trimming primarily reflected the removal of reads that became shorter than the minimum accepted length, rather than poor sequencing quality, providing a transparent explanation for read loss during small RNA-seq preprocessing. Using this framework, we identified a small set of high-confidence *de novo* miRNA candidates and integrated them with a confounding-corrected transcriptomic signature, revealing regulatory networks associated with inflammation, extracellular matrix remodeling, autophagy, and cellular adaptation. Importantly, although based on publicly available datasets, this study focuses exclusively on female cohorts, contributing to ongoing efforts to reduce the persistent underrepresentation of women in molecular exercise research. A large-scale analysis of the exercise science literature reported that women represent only approximately one-third of study participants, highlighting the need for analytical frameworks applicable to female populations ^43^.

Comprehensive seed-space analysis revealed that five of the seven final candidates possessed previously unreported canonical seed sequences, whereas two shared canonical seeds with annotated human miRNA families. Additional non-canonical 7-mer similarities were identified elsewhere within the mature sequences, including matches to members of the miR-16/195 family. These similarities occur outside the canonical seed region (positions 2–8) and therefore do not imply conservation of the primary target-recognition motif. Instead, they indicate that portions of the mature sequence overlap with known miRNA seed repertoires, suggesting partial sequence conservation without necessarily implying shared regulatory targets. Interestingly, members of the miR-16 family have been associated with the systemic somatic stress response to exercise (Supplementary Table 7), although whether these partial sequence similarities reflect functional convergence remains to be experimentally determined.

It should also be noted that each *de novo* candidate represents a CD-HIT cluster rather than a single unique sequence. Because CD-HIT selects cluster representatives using a greedy algorithm, the representative sequence provides a reproducible reference for downstream analyses but is not necessarily the most abundant or biologically relevant isoform ^44^. Consequently, alternative cluster members may also represent biologically relevant sequence variants, an aspect that warrants future investigation in experimental datasets specifically designed to characterize isomiR diversity.

Integration with RNA-seq data further demonstrated that biologically meaningful regulatory relationships can be recovered despite substantial tissue-driven variability. After accounting for major confounding effects, the resulting transcriptomic signature remained enriched for immune activation, cytotoxic functions, metabolic regulation, and cellular homeostasis, consistent with previous observations in exercise biology ^45^; ^37^; ^46^. Integrating these genes with predicted miRNA targets expanded the reconstructed regulatory network, revealing convergence of both *de novo* and known miRNAs on pathways involved in extracellular matrix organization, autophagy, inflammation, and muscle remodeling. The complementary validation of predicted miRNA–mRNA interactions using multiMiR and miRanda further increased confidence in these regulatory associations. As an example of its downstream applications, we further interrogated a curated set of genes associated with sarcopenia, a major skeletal muscle disorder, and identified validated interactions involving pathways related to inflammation, muscle atrophy, extracellular matrix remodeling, mitochondrial metabolism, senescence, and myogenesis. This targeted analysis illustrates how nf-sarcopipe can be readily applied to prioritize disease-relevant regulatory hypotheses from broader transcriptomic datasets.

This study has several limitations. The integrative analysis was performed using independent sRNA-seq and RNA-seq datasets generated from different biological tissues, requiring statistical correction to minimize tissue-driven confounding. Consequently, the identified miRNA–mRNA interactions should be interpreted as biologically coherent associations rather than direct evidence of intracellular regulation. In addition, the *de novo* candidates identified here remain computational predictions that require experimental validation to confirm their expression, processing, and regulatory activity.

Future studies using matched sRNA-seq and RNA-seq datasets generated from the same individuals and biological compartments will enable more direct characterization of miRNA–mRNA regulatory interactions while providing an opportunity to experimentally validate the novel candidates identified in this work. Together, these results demonstrate that nf-sarcopipe provides a robust computational framework for integrating *de novo* and reference-guided miRNA discovery with transcriptomic data, facilitating the identification of biologically meaningful regulatory networks associated with exercise adaptation, muscle remodeling, and healthy aging.

## Conclusion

Overall, nf-sarcopipe provides a scalable and reproducible end-to-end pipeline that unifies *de novo and* reference-guided miRNA discovery with transcriptome-wide integration, differential expression analysis, structural filtering of *de novo* candidates, seed-based target prediction, and complementary validation of miRNA–mRNA interactions using multiMiR and miRanda. This integrated framework enables the systematic prioritization of high-confidence regulatory miRNAs and their putative targets, facilitating downstream functional interpretation of exercise-associated molecular adaptations.

## Materials and Methods

### 1. - miRNA and mRNA dataset

Small RNA sequencing (sRNA-seq) data were obtained from a publicly available plasma dataset including healthy young synchronized swimmers (Active, n = 10) and sedentary women (Sedentary, n = 15) reported by ^47^. For transcriptomic analyses, publicly available RNA-seq datasets from healthy young physically active female skaters (whole blood, n = 7;^48^) and sedentary women (vastus lateralis muscle biopsies, n = 6;^49^) were used. Because the RNA-seq datasets originated from different tissues and independent cohorts, additional analyses were performed to assess and correct tissue-driven confounding before downstream miRNA–mRNA integration.

### 2. - nf-sarcopipe implementation

The pipeline was implemented in Nextflow v25.10.2 using the DSL2 framework to provide a modular, scalable, and reproducible pipeline for the analysis and integration of small RNA sequencing (sRNA-seq) and mRNA sequencing datasets. nf-sarcopipe is organized into three main modules: (1) Preprocessing, (2) miRNA Discovery, and (3) Target Prediction & mRNA Integration. The first two modules process sRNA-seq data to identify both reference-guided and *de novo* miRNAs, whereas the third module integrates these predictions with differential gene expression results obtained from RNA-seq data processed using the standardized nf-core/rnaseq workflow (v3.22.2) ^50^. This integration enables transcriptome-wide target prediction, coherent miRNA–mRNA interaction analysis, and generation of network-ready outputs for downstream biological interpretation. Each module consists of independent processes connected through Nextflow channels, enabling efficient parallel execution across local and high-performance computing (HPC) environments. All software dependencies were managed through containerized environments to ensure computational reproducibility. Pipeline execution was automatically documented through Nextflow execution reports, workflow traces, directed acyclic graphs (DAGs), and resource usage summaries, including CPU utilization, memory consumption, and execution time (15 h 45 min). The workflow supports both local execution and SLURM-managed HPC systems.

#### a) Small RNA-seq filtering

Raw small RNA sequencing reads were first subjected to quality control and adapter removal. Adapter trimming and quality filtering were performed using fastp v0.23.4 ^51^ and the adapter sequences provided by the sequencing facility, ensuring the removal of library-specific adapter contamination while preserving short reads with lengths consistent with small RNA libraries (18–28 nt).

Quality-filtered reads were then processed for downstream miRNA discovery and reference-guided identification. All preprocessing steps were implemented within the sarcopipe preprocessing module to ensure standardized input data for subsequent analyses.

#### b) miRNA identification (BrumiR and miRDeep2)

miRNA candidates were identified using two complementary algorithms. *De novo* miRNA discovery was performed using BrumiR v3.0, which identifies candidate miRNAs directly from small RNA sequencing data without requiring a reference genome. To increase confidence in candidate identification, BrumiR outputs were subsequently filtered using the BrumiR-RF module, a machine-learning classifier trained on experimentally validated miRNAs. Unlike the initial *de novo* prediction performed by BrumiR, this filtering step uses the human reference genome (GRCh38) to evaluate and retain high-confidence putative miRNA candidates.

Reference-guided miRNA identification was performed using miRDeep2 v2.0.1.3. In this branch, reads were mapped to the human reference genome GRCh38 using the mapper module implemented in miRDeep2, and known miRNAs were identified through alignment to reference miRNA databases. This dual-algorithm approach enabled the detection of both known and novel miRNA candidates.

### 3. - Module miRNA Discovery

#### a) Candidate clustering and expression matrix construction

To reduce redundancy among miRNA candidates, sequences identified by BrumiR were clustered using CD-HIT-est v4.8.1 ^52^ based on sequence similarity. Multiple clustering identity thresholds were evaluated (85%, 90%, and 95%), and the 95% identity threshold was selected for all downstream analyses. Representative sequences from each cluster were retained, and cluster membership information was subsequently used to construct presence–absence tables across samples.

Expression matrices were generated for both BrumiR candidates and miRDeep2-identified miRNAs by aggregating read counts across samples. To increase robustness and reduce spurious detections, sequences predicted by each algorithm were retained only if present in at least 80% of the samples within each experimental group (presence threshold p = 0.8). These filtered matrices provided the basis for subsequent comparative analyses and differential expression testing.

#### b) Comparative analysis of BrumiR and miRDeep2 predictions

To evaluate the overlap between miRNA candidates identified by BrumiR and those detected by miRDeep2, sequence-level comparisons were performed. Initial overlap analyses were based on exact sequence matches between both datasets.

To further assess biological concordance and annotation status, miRNA candidates were aligned against the curated miRGeneDB reference database using BLAST (blastn-short) ^53,54^. Stringent similarity thresholds were applied (≥98% sequence identity and ≥75% query coverage, effectively restricting retained alignments to perfect matches for mature miRNA sequences) to identify high-confidence matches while allowing for minor sequence variation associated with isomiRs and end-processing heterogeneity ^55^. Sequence identity corresponds to the proportion of identical nucleotides between the query and reference sequences, whereas query coverage reflects the fraction of the query sequence aligned to the reference. The selected thresholds balance sensitivity and specificity for short RNA sequences, ensuring robust identification of conserved miRNAs while minimizing false-positive annotations.

Core miRNA sets were defined based on their presence across samples within each experimental group. Comparative analyses were performed to quantify overlap between algorithms and to evaluate the consistency of miRNA detection across datasets, distinguishing shared annotated miRNAs from putative novel candidates.

### 4. - Module Target Prediction & mRNA Integration

#### a) Structural support analysis of de novo miRNA candidates

Candidate miRNAs identified *de novo* by Brumir-RF were further evaluated for structural support by mapping candidate sequences to the reference genome (BrumiR2Reference branch) and extracting the corresponding precursor regions. RNA secondary structures were predicted using RNAfold from the ViennaRNA package v2.7.2.

Predicted hairpin structures and associated minimum free energy values were used to assess the structural plausibility of candidate miRNAs. Structural support metrics were compiled into summary tables to facilitate downstream evaluation of candidate reliability.

#### b) miRNA differential expression analysis

Expression matrices generated for both BrumiR candidates and miRDeep2-identified miRNAs were analyzed for differential expression between Active and Sedentary groups. Differential expression analysis was performed in R v4.5.0 using the DESeq2 v1.48.2 ^56^ framework. Raw counts were normalized using the median-of-ratios method, and statistical significance was evaluated using the Wald test with Benjamini–Hochberg correction for multiple testing.

miRNAs with adjusted p-values below 0.05 were considered significantly differentially expressed and were retained for downstream integrative analyses.

#### c) Sequence-based annotation and overlap analysis of miRNA candidates

To assess the degree of similarity between miRNA candidates identified by BrumiR-RF and miRDeep2 (output from miRNA Discovery module), as well as their correspondence with annotated miRNAs, sequence-level comparisons were performed. Mature miRNA sequences were aligned against curated reference databases using BLASTN v2.12.0+ (NCBI BLAST+ suite) ^57^ with parameters optimized for short RNA sequences (`-task blastn-short`). Stringent alignment thresholds were applied, including a minimum sequence identity of 98% sequence identity and ≥75% query coverage (effectively restricting retained alignments to perfect matches for mature miRNA sequences), to ensure high-confidence matches while minimizing false-positive annotations.

Reference datasets included miRGeneDB (human mature sequences) ^58^ for high-confidence annotation and miRBase for complementary sequence exploration ^34^. Overlap analyses were conducted by comparing BrumiR-RF high-confidence candidates and miRDeep2-derived miRNAs both at the sequence level (exact matches) and at the annotation level (shared database hits). The resulting intersections were used to quantify shared and unique miRNA candidates across methods and to distinguish putative novel miRNA candidates from known miRNAs.

#### d) 7-mer seed-space analysis of miRNA candidates

To further characterize the novelty of BrumiR-derived miRNA candidates beyond canonical seed definitions, a comprehensive 7-mer seed-space analysis was performed ^59^. For each mature miRNA sequence, all possible contiguous 7-mer subsequences were generated by sliding a window across the full length of the sequence.

A reference dictionary of canonical miRNA seeds was constructed from annotated human miRNAs obtained from miRBase, extracting the seed region corresponding to nucleotides 2–8 of each mature sequence. Each candidate-derived 7-mer was then compared against this reference set to identify matches to known seed sequences.

For each candidate, the following metrics were computed: total number of 7-mers, number of 7-mers matching known seeds, positional distribution of matches, and associated miRNA families. The canonical seed region (positions 2–8) was separately classified as either known or novel. Two control sequences were included to validate the analysis: a known miRNA (hsa-miR-660-5p) and a sequence containing a known seed motif (known_seed_A).

This analysis allowed the identification of candidates with complete absence of similarity to known miRNA seeds, as well as those exhibiting partial similarity at non-canonical positions. The approach provides a complementary layer of sequence-based characterization, enabling discrimination between fully novel candidates and sequences potentially related to known miRNA families through seed similarity outside the canonical region.

#### e) mRNA-seq processing, confounding correction and integrative analysis

RNA-seq datasets were processed using the nf-core/rnaseq pipeline (v3.22.2) implemented in Nextflow, which performs quality control, adapter trimming, genome alignment, transcript quantification, and generation of gene-level count matrices. Raw paired-end reads generated by GEPREP v3.2 ^60^ were assessed using FastQC before and after trimming to verify sequencing quality.

Lowly expressed genes were filtered by retaining transcripts with at least 10 counts in a minimum of two samples. Differential expression analysis was performed in R (v4.5.0) using DESeq2 (v1.48.2). Count normalization was carried out using the median-of-ratios method, and variance-stabilizing transformation (VST) was applied for exploratory analyses and data visualization.

Because the RNA-seq datasets originated from distinct tissues (whole blood and skeletal muscle), principal component analysis (PCA) was performed on the 5,000 most variable genes to identify major sources of variation ^61^ ^62^.

Gene expression was subsequently modeled using linear regression with principal components included as covariates to account for tissue-driven confounding. Based on sensitivity analyses, the final model included PC1 and PC2:

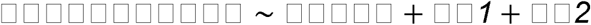

where *Expression_g_* represents the expression level of gene *g*, Group corresponds to the biological condition (Active vs. Sedentary), and PC1 and PC2 were included as covariates to account for tissue-driven confounding.

Statistical significance was assessed using Benjamini–Hochberg false discovery rate (FDR) correction, and genes with adjusted *p* < 0.05 were considered differentially expressed.

To further evaluate the contribution of cellular composition, immune deconvolution was performed on whole-blood samples using quanTIseq implemented in the immunedeconv framework after TPM normalization and HGNC gene symbol annotation.

Finally, the confounding-corrected differentially expressed genes were integrated with circulating miRNA profiles for downstream miRNA–mRNA regulatory analyses. Experimentally supported genes and computationally predicted targets were analyzed separately to distinguish high-confidence observations from hypothesis-generating regulatory candidates.

#### f) miRNA–mRNA integration and regulatory network reconstruction

Putative miRNA–mRNA regulatory interactions were inferred by integrating differentially expressed miRNAs with confounding-corrected RNA-seq results. Canonical seed regions (positions 2–8) were extracted from both BrumiR-derived *de novo* miRNAs and miRDeep2-identified known miRNAs. Reverse-complement seed sequences were systematically scanned against 3′ untranslated regions (3′UTRs) using exact 7-mer seed matching to identify candidate target transcripts^59^.

Predicted interactions were filtered according to expression coherence, retaining only inversely regulated miRNA–mRNA pairs that were consistent with the canonical repressive mode of miRNA action. Accordingly, interactions were retained when an upregulated miRNA was paired with a downregulated target gene or, conversely, when a downregulated miRNA was paired with an upregulated target gene ^59,63^. For known miRNAs identified by miRDeep2, target predictions were retrieved using the multiMiR R package, which integrates experimentally validated and computationally predicted miRNA–target interactions from multiple public databases ^64^. For BrumiR-derived *de novo* candidates, target predictions were independently evaluated using miRanda, applying sequence complementarity and duplex stability criteria to support candidate regulatory interactions ^65^.

Coherent interactions obtained from both approaches were merged into a unified set of candidate miRNA–mRNA regulatory pairs. Gene-level target lists were generated separately for BrumiR and miRDeep2 and combined to produce a non-redundant union for downstream functional analyses.

For network reconstruction, all coherent interactions were compiled into Cytoscape-compatible edge and node tables. Node attributes included molecule type (miRNA or gene), miRNA origin (BrumiR or miRDeep2), differential expression status, and regulatory direction, whereas edges represented predicted regulatory interactions supported by seed matching together with miRanda (*de novo* candidates) or multiMiR (known miRNAs). The resulting networks were visualized using Cytoscape v3.10.3 ^66^ to facilitate downstream exploration of candidate regulatory modules associated with exercise adaptation and skeletal muscle biology.

## Availability of Supporting Source Code and Requirements

Project name: sarcopipe

Project home page: https://github.com/AGENslab/nf-sarcopipe

Manuscript analysis scripts: https://github.com/AGENslab/nf-sarcopipe_ms

Operating system(s): Unix and Linux (tested on high-performance computing clusters using SLURM workload manager)

Programming language: Nextflow, Python, AWK, and Shell

Other requirements: Nextflow DSL2, fastp, BrumiR, BrumiR-RF, BrumiR2Reference, miRDeep2, Bowtie (v1), CD-HIT-est and CD-HIT-est-2d, Python 3, and R (with DESeq2) for downstream analyses.

Schematic illustrations: Created with BioRender.com License: MIT License

Any restrictions to use by nonacademics: none

## Additional Files

Supplementary Figure 1. Robustness of de novo candidate detection and evaluation of miRDeep2 novel predictions.

Supplementary Figure 2. Structural and sequence-based characterization of de novo miRNA candidates identified by nf-Sarcopipe.

Supplementary Figure 3. Evidence of confounding effects and their correction in RNA-seq analysis.

Supplementary Figure 4. Extended analysis of predicted miRNA–mRNA interactions and pathway enrichment.

Supplementary Table 1. Sample metadata and preprocessing summary for the plasma sRNA-seq dataset used for initial small RNA candidate discovery.

Supplementary Table 2 (page 1 & 2). Summary of raw library quality and fastp filtering statistics across the 25 small RNA-seq libraries.

Supplementary Table 3. Unique structurally supported BrumiR-RF core candidate miRNAs identified after BrumiR2Reference analysis.

Supplementary Table 4. Characteristics of the final high-confidence de novo miRNA candidates identified by nf-Sarcopipe.

Supplementary Table 5. Metadata and sequencing quality metrics for the RNA-seq datasets used in the integrative analysis.

Supplementary Table 6. Functional classification of sarcopenia-associated genes.

Supplementary Table 7 (page 1&2). Curated list of microRNAs associated with exercise responses in humans.

## Competing Interests

The authors declare that they have no competing interests.

## Funding

This work was supported by the interdisciplinary research grant Fondos Inter UOH 005-2024 from Universidad de O’Higgins, and by the Agencia Nacional de Investigación y Desarrollo (ANID) through FONDECYT grants 1241959 (D.V.I.) and 11251927 (C.M).

## Authors Contributions

N.P.D. designed, developed, and implemented the Sarcopipe workflow, performed the computational analyses, and wrote the initial version of the manuscript. C.M guided the development of nf-sarcopipe. F.G.M. and G.C.M. contributed to the structuring, technical revision, and validation of the pipeline for repository release. C.M. and A.D.G. contributed to methodological discussion, provided technical guidance, and assisted in the resolution of complex analytical challenges. D.V.I provided crucial biological feedback. C.M. provided crucial computational feedback. C. M and D.V.I helped to improve the manuscript. C.M and D.V.I. supervised the study and contributed to manuscript revision. All authors provided helpful discussions for the work and reviewed the manuscript. All authors approved the final manuscript.

## Acknowledgements

This research was performed using the Kütral computer cluster at the Computational Biology Laboratory (CBLab), Universidad de O’Higgins, Rancagua, Chile. Kütral is a high-performance computing infrastructure designed for large-scale genomics analyses (funding FIC40059065).

## Contributor Information

Natalia Poblete-Durán, Instituto de Ciencias de la Salud; Instituto de Ciencias de la Ingeniería, Universidad de O’Higgins, 2820000 Rancagua, Chile.

Felipe Gómez-Molina, Instituto de Ciencias de la Ingeniería, Universidad de O’Higgins, 2820000 Rancagua, Chile.

Gabriel Cabas-Mora, Instituto de Ciencias de la Ingeniería, Universidad de O’Higgins, 2820000 Rancagua, Chile.

Alex Di Genova-Bravo, Instituto de Ciencias de la Ingeniería, Universidad de O’Higgins, 2820000 Rancagua, Chile.

Denisse Valladares-Ide, Instituto de Ciencias de la Salud, Universidad de O’Higgins, 2820000 Rancagua, Chile.

Carol Moraga-Quinteros, Instituto de Ciencias de la Ingeniería, Universidad de O’Higgins, 2820000 Rancagua, Chile.

## Table 6 references

67–78

## Table 7 references

79–108

## Notes

https://github.com/AGENslab/nf-sarcopipe

### Competing Interest Statement

The authors have declared no competing interest.

