## Supplementary Figures for "nf-sarcopipe enables integrative discovery of exercise-responsive miRNAs and miRNA–mRNA regulatory networks associated with skeletal muscle adaptation"

†Corresponding authors:

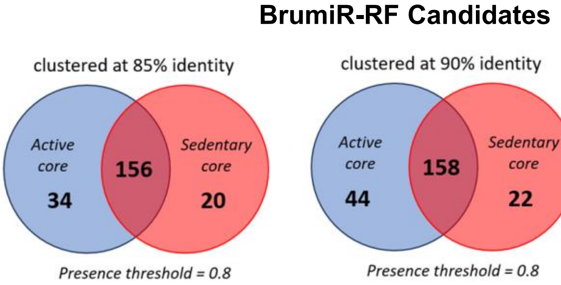


| CD-HIT identity | Total clusters | Singletons n (%) | Ubiquitous (25/25) n (%) | Weighted median presence | Active core | Sedentary core | Shared core | Core union |
|---|---|---|---|---|---|---|---|---|
| 0.85 | 836 | 360 (43.1%) | 95 (11.4%) | 2 | 190 | 176 | 156 | 210 |
| 0.90 | 1028 | 421 (41.0%) | 95 (9.2%) | 3 | 202 | 180 | 158 | 224 |
| 0.95 | 1307 | 542 (41.5%) | 94 (7.2%) | 2 | 198 | 174 | 152 | 220 |

b)

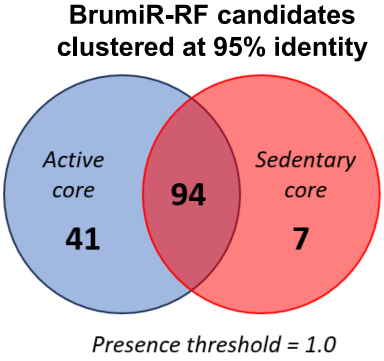


c)

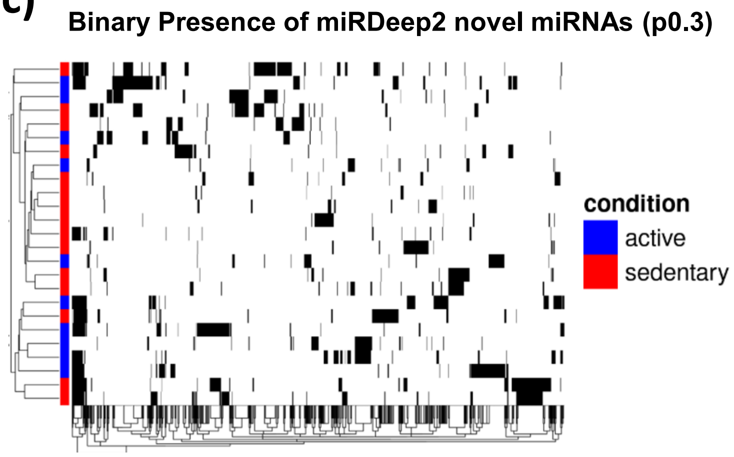


d)

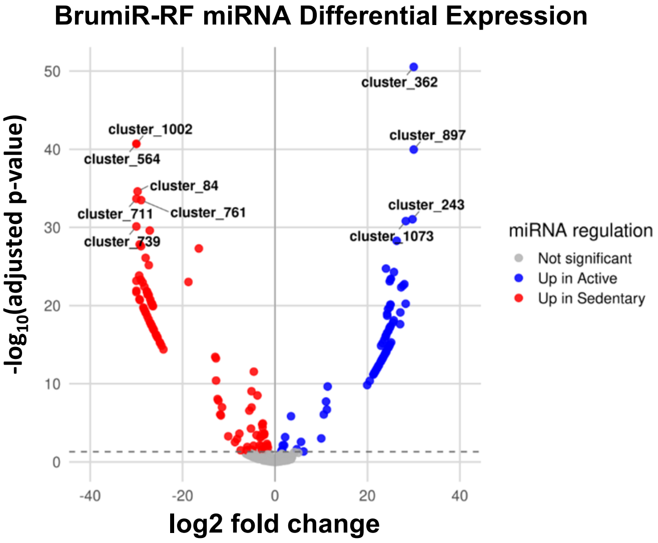


e)

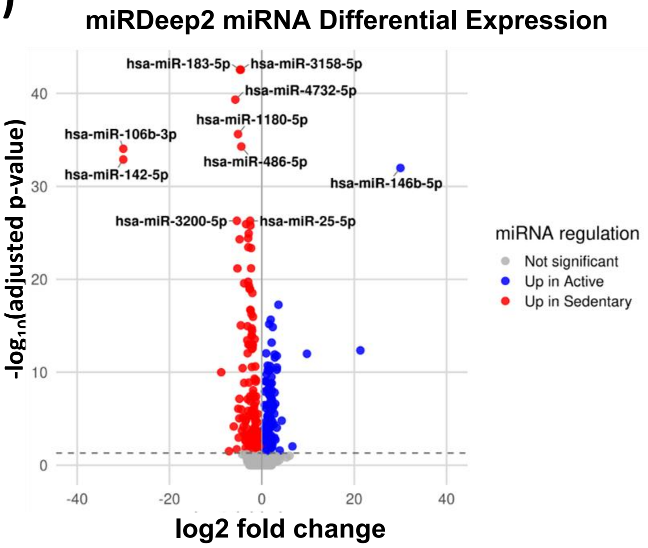

**Supplementary Figure 1.** ***Robustness of de novo candidate detection and evaluation of miRDeep2 novel predictions.*** (a) BrumiR-RF candidate miRNAs clustered at 85%, 90% and 95% sequence identity showing the overlap between active and sedentary core sets (presence ≥0.8). Increasing clustering stringency increases the total number of clusters while maintaining stable core set composition across conditions. Summary statistics include total clusters, proportion of singletons, ubiquitous clusters (detected in all samples), and core set sizes. (b) Core miRNA candidates identified by BrumiR-RF at 95% identity using a stringent presence threshold (≥1.0), showing reduced set size compared to the ≥0.8 threshold while preserving all of shared candidates.(c) Binary presence heatmap of miRDeep2 novel miRNAs detected at a relaxed presence threshold (≥0.3). The sparse and heterogeneous distribution of these candidates across samples indicates limited reproducibility compared to core miRNA sets used in downstream analyses. (d–e) Volcano plots showing differential expression results for BrumiR-RF candidates (d) and miRDeep2 known miRNAs highlighting the distribution of effect sizes and statistical significance across methods.

a)

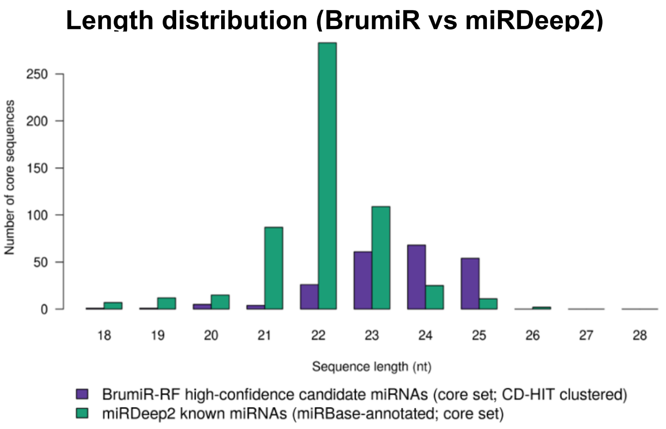


b)

**Cluster size of BrumiR passfilter *de novo* miRNA candidates**

| Final candidate | Original cluster | Group | Cluster size | Classification |
|---|---|---|---|---|
| hsa-miR-novel_A | cluster_740 | active | 9 | Cluster |
| hsa-miR-novel_B | cluster_986 | active | 11 | Cluster |
| hsa-miR-novel_C | cluster_42 | shared | 27 | Cluster |
| hsa-miR-novel_D | cluster_374 | shared | 29 | Cluster |
| hsa-miR-novel_E | cluster_413 | shared | 26 | Cluster |
| hsa-miR-novel_F | cluster_465 | shared | 32 | Cluster |
| hsa-miR-novel_G | cluster_485 | shared | 23 | Cluster |

c)

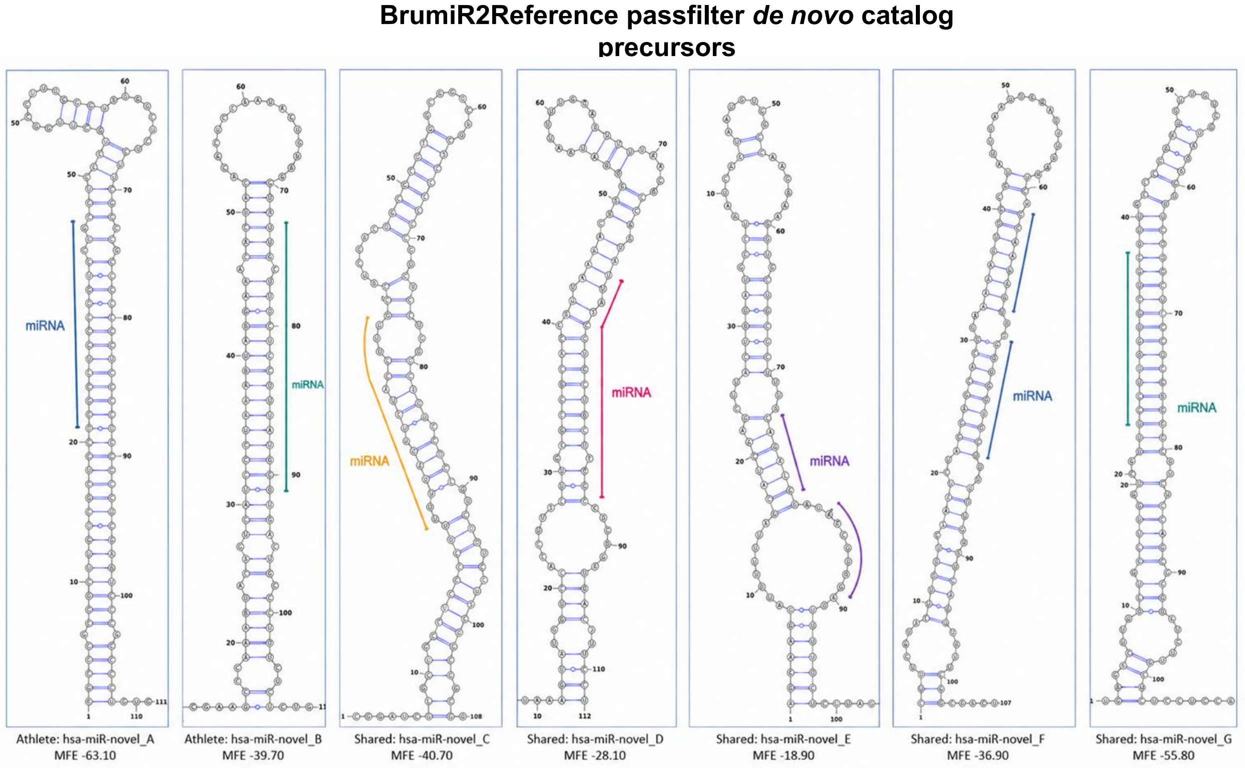


d)

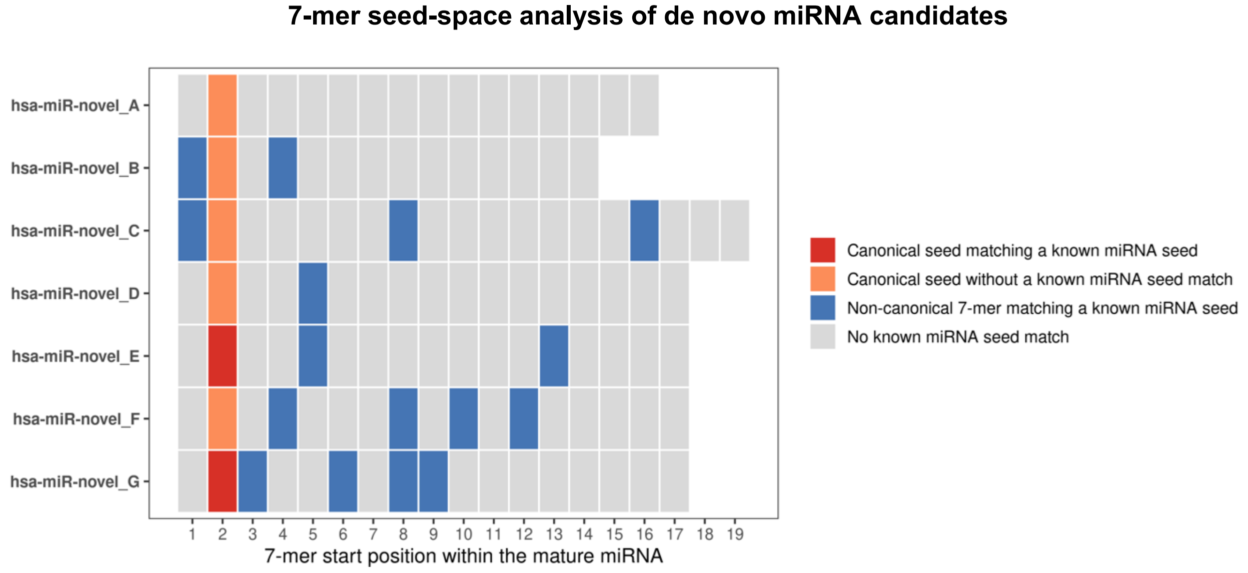

**Supplementary Figure 2.** ***Structural and sequence-based characterization of de novo miRNA candidates identified by nf-Sarcopipe***. (a) Length distribution of miRNA candidates detected by BrumiR-RF and miRDeep2. BrumiR high-confidence *de novo* candidates show a broader size distribution enriched at 22–25 nucleotides, whereas miRDeep2-known miRNAs display a peak at 22 nucleotides, consistent with canonical Dicer processing. (b) Summary of the final high-confidence *de novo* miRNA candidates retained after BrumiR2Reference structural validation. For each candidate, the original CD-HIT cluster, group specificity, cluster size, and candidate classification are shown. Cluster size corresponds to the number of sequences grouped within each representative CD-HIT cluster. (c) Predicted secondary structures of the 7 precursor sequences for the BrumiR2Reference passfilter *de novo* miRNA candidates visualized using VARNA (red dots indicate miRNA site). Mature miRNA regions are highlighted within the stem region of the hairpin structures, and MFE values indicate stable folding compatible with canonical miRNA precursors. (d) 7-mer seed-space analysis of BrumiR passfilter candidates miRNAs. All possible 7-mer subsequences derived from mature miRNA sequences were compared against canonical human miRNA seeds from miRBase. The heatmap shows the presence or absence of matches across all possible 7-mer positions within each sequence. The canonical seed region (nucleotides 2–8) is highlighted and classified as either known or novel.

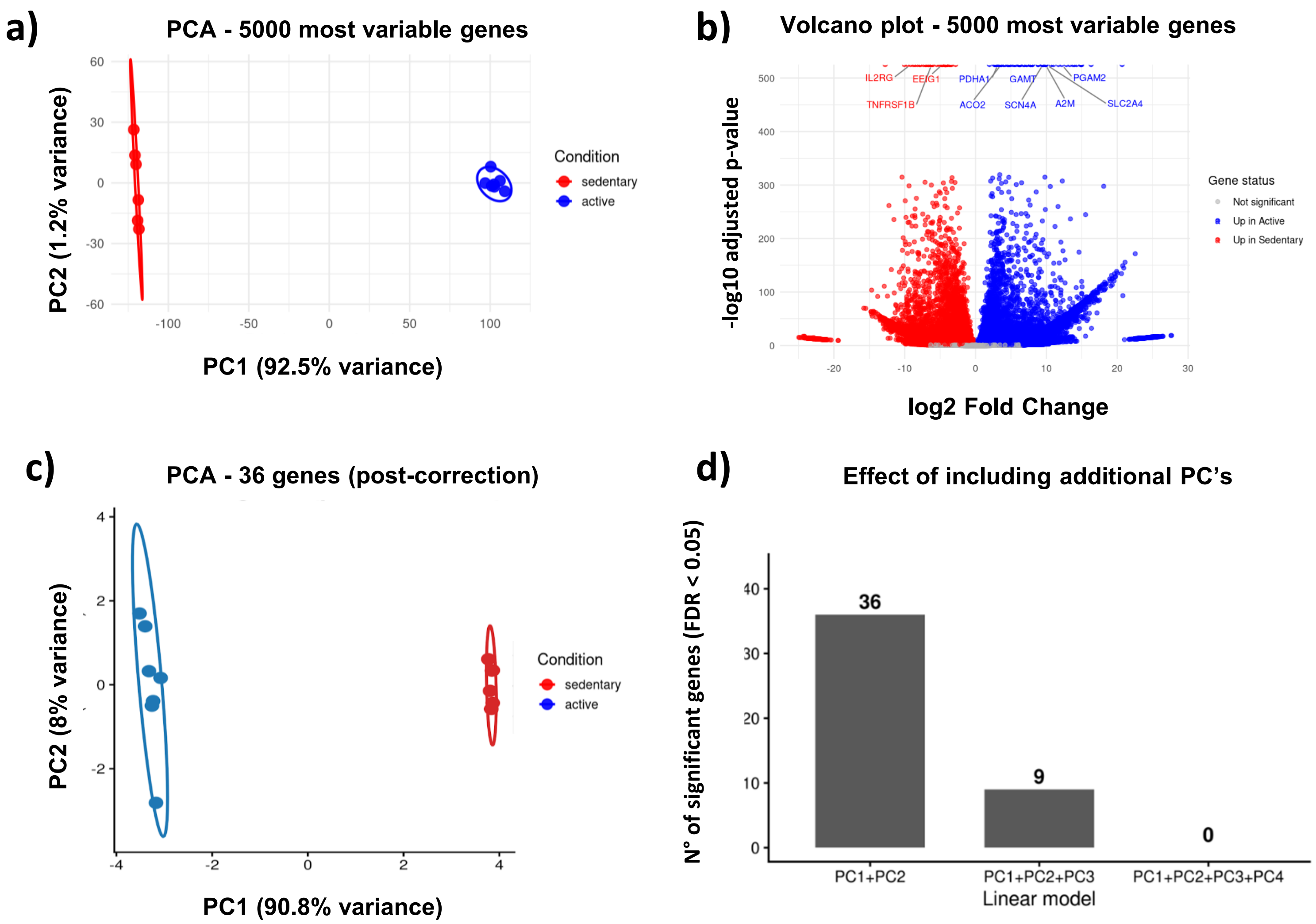
a)
PCA - 5000 most variable genes
PC2 (1.2% variance)
PC1 (92.5% variance)
Condition
sedentary
active
b)
Volcano plot - 5000 most variable genes
-log10 adjusted p-value
log2 Fold Change
IL2RG
EEIG1
TNFRSF1B
PDHA1
GAMT
PGAM2
ACO2
SCN4A
A2M
SLC2A4
Gene status
Not significant
Up in Active
Up in Sedentary
c)
PCA - 36 genes (post-correction)
PC2 (8% variance)
PC1 (90.8% variance)
Condition
sedentary
active
d)
Effect of including additional PC's
N° of significant genes (FDR < 0.05)
36
9
0
PC1+PC2
PC1+PC2+PC3
PC1+PC2+PC3+PC4
Linear model

**Supplementary Figure 3.** ***Evidence of confounding effects and their correction in RNA-seq analysis.*** (a) Principal component analysis (PCA) of the 5,000 most variable genes showing strong separation between samples driven by tissue origin, with PC1 explaining 92.5% of the variance. (b) Volcano plot of differential expression analysis without correction, revealing a large number of significant genes consistent with a confounded signal dominated by tissue-specific effects. (c) PCA of the 36 genes retained after correction (expression ~ group + PC1 + PC2), showing separation of samples according to biological condition rather than tissue origin, indicating effective removal of the dominant confounding effect. (d) Sensitivity analysis showing the effect of including additional principal components in the linear model. The number of significantly differentially expressed genes (FDR < 0.05) decreases from 36 (PC1 + PC2) to 9 (PC1 + PC2 + PC3) and 0 (PC1 + PC2 + PC3 + PC4), demonstrating over-adjustment and loss of biological signal when higher-order components are included.

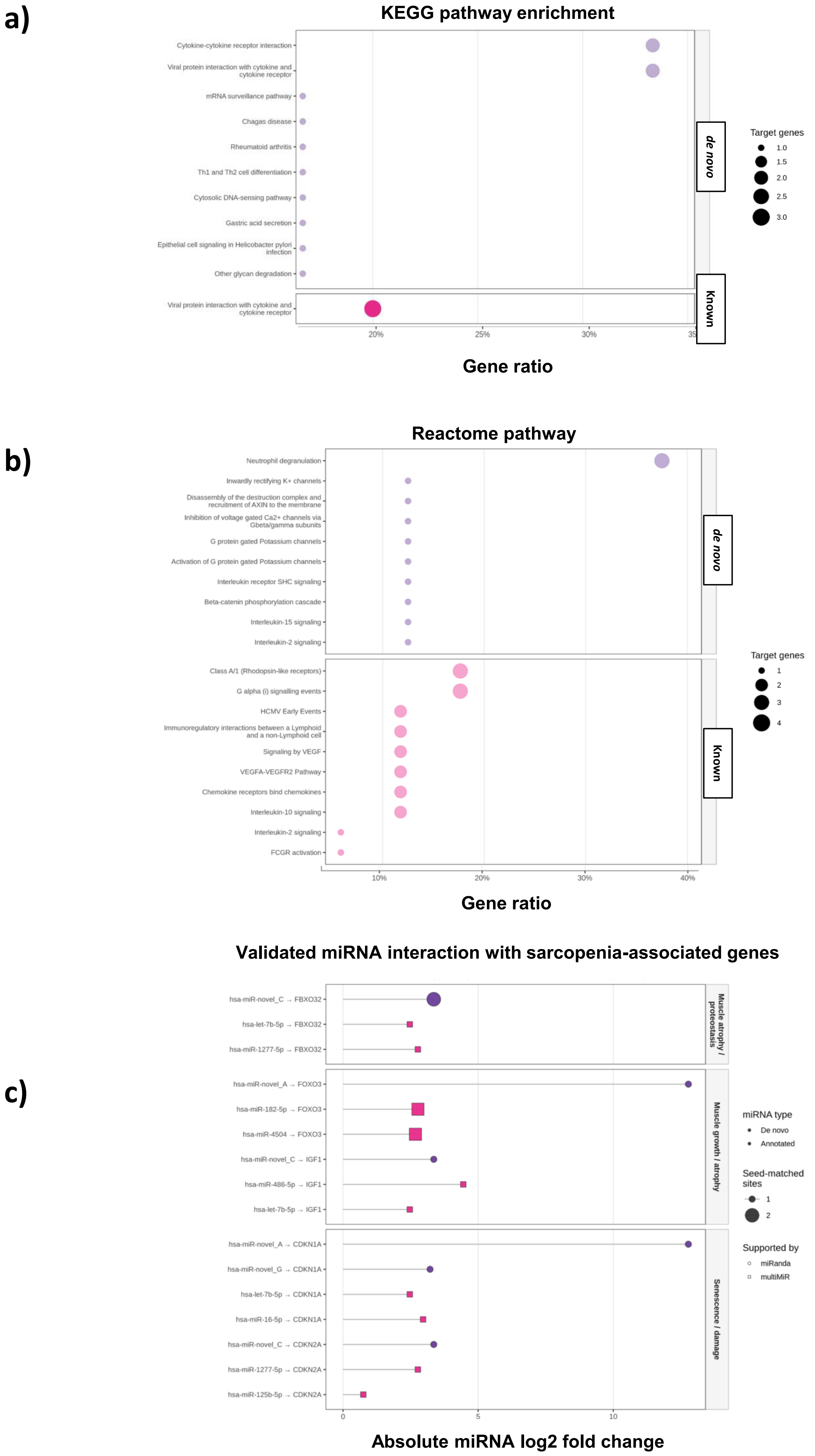

a)
KEGG pathway enrichment
Cytokine-cytokine receptor interaction
Viral protein interaction with cytokine and cytokine receptor
mRNA surveillance pathway
Chagas disease
Rheumatoid arthritis
Th1 and Th2 cell differentiation
Cytosolic DNA-sensing pathway
Gastric acid secretion
Epithelial cell signaling in Helicobacter pylori infection
Other glycan degradation
Viral protein interaction with cytokine and cytokine receptor
de novo
Known
20%
25%
30%
Gene ratio
Target genes
1.0
1.5
2.0
2.5
3.0
b)
Reactome pathway
Neutrophil degranulation
Inwardly rectifying K+ channels
Disassembly of the destruction complex and recruitment of AXIN to the membrane
Inhibition of voltage gated Ca2+ channels via Gbeta/gamma subunits
G protein gated Potassium channels
Activation of G protein gated Potassium channels
Interleukin receptor SHC signaling
Beta-catenin phosphorylation cascade
Interleukin-15 signaling
Interleukin-2 signaling
Class A/1 (Rhodopsin-like receptors)
G alpha (i) signalling events
HCMV Early Events
Immunoregulatory interactions between a Lymphoid and a non-Lymphoid cell
Signaling by VEGF
VEGFA-VEGFR2 Pathway
Chemokine receptors bind chemokines
Interleukin-10 signaling
Interleukin-2 signaling
FCGR activation
de novo
Known
10%
20%
30%
40%
Gene ratio
Target genes
1
2
3
4
c)
Validated miRNA interaction with sarcopenia-associated genes
hsa-miR-novel_C → FBXO32
hsa-let-7b-5p → FBXO32
hsa-miR-1277-5p → FBXO32
hsa-miR-novel_A → FOXO3
hsa-miR-182-5p → FOXO3
hsa-miR-4504 → FOXO3
hsa-miR-novel_C → IGF1
hsa-miR-486-5p → IGF1
hsa-let-7b-5p → IGF1
hsa-miR-novel_A → CDKN1A
hsa-miR-novel_G → CDKN1A
hsa-let-7b-5p → CDKN1A
hsa-miR-16-5p → CDKN1A
hsa-miR-novel_C → CDKN2A
hsa-miR-1277-5p → CDKN2A
hsa-miR-125b-5p → CDKN2A
Muscle atrophy / proteostasis
Muscle growth / atrophy
Senescence / damage
0
5
10
Absolute miRNA log2 fold change
miRNA type
De novo
Annotated
Seed-matched sites
1
2
Supported by
miRanda
multiMiR

**Supplementary Figure 4. *Extended analysis of predicted miRNA–mRNA interactions and pathway enrichment*.** (a) KEGG pathway enrichment analysis performed separately on validated target genes regulated by *de novo* and annotated miRNAs. The *de novo* miRNA targets were associated with several immune- and signaling-related pathways that did not remain significant after multiple-testing correction, whereas the annotated miRNA target set showed significant enrichment for the cytokine–cytokine receptor interaction pathway. (b) Reactome pathway enrichment analysis performed separately for validated targets of *de novo* and annotated miRNAs. Although no pathways remained significant after multiple-testing correction, the highest-ranking Reactome terms for *de novo* miRNA targets included interleukin-2/interleukin-15 signaling, G protein-coupled receptor signaling, β-catenin-related processes, and neutrophil degranulation, supporting the biological plausibility of the prioritized candidates. (c) Validation of predicted interactions involving a curated set of musculoskeletal genes associated with muscle growth, atrophy, extracellular matrix remodeling, mitochondrial metabolism, inflammation, and cellular senescence. Only interactions supported by miRanda (*de novo* miRNAs) or multiMiR (annotated miRNAs) are shown. Point size represents the number of canonical seed-matched sites, whereas color and shape distinguish the validation strategy and miRNA discovery branch.

| ID | Sex | Excercise | State | Health Status | miRNA Biopsy | Reference | raw reads | Proceced reads (fastp + miRNA adapters [small RNA-seq]) | Proceced reads (fastp + miRNA adapters + splitter) | | Brumir-core Short | | | Brumir-RF | miRDeep2 Short | |
|---|---|---|---|---|---|---|---|---|---|---|---|---|---|---|---|---|
| | | | | | | | | | Reads between 18–28 nt (short reads) | >28 | UNITIGS | Candidates | Other sequence | High-conffidence miRNAs | miRNAs novel | miRNAs known |
| SRR11363960 | Female | no | Sedentary 10 | Healthy | Plasma | Li et al. 2020 | 19.022.725 | 9.845.947 | 9.084.749 | 761.198 | 61.668 | 615 | 834 | 402 | 1.378 | 850 |
| SRR11363961 | Female | no | Sedentary 9 | Healthy | Plasma | Li et al. 2020 | 20.590.468 | 8.501.859 | 8.136.652 | 365.207 | 28.873 | 484 | 236 | 365 | 1.214 | 804 |
| SRR11363962 | Female | no | Sedentary 8 | Healthy | Plasma | Li et al. 2020 | 23.587.262 | 13.814.661 | 13.071.752 | 742.909 | 63.781 | 691 | 504 | 480 | 1.422 | 755 |
| SRR11363963 | Female | no | Sedentary 7 | Healthy | Plasma | Li et al. 2020 | 20.654.567 | 6.128.209 | 5.577.336 | 550.873 | 74.782 | 547 | 690 | 360 | 1.219 | 788 |
| SRR11363964 | Female | no | Sedentary 6 | Healthy | Plasma | Li et al. 2020 | 23.241.588 | 2.871.766 | 2.258.482 | 613.284 | 88.711 | 348 | 950 | 224 | 907 | 647 |
| SRR11363965 | Female | no | Sedentary 5 | Healthy | Plasma | Li et al. 2020 | 20.028.716 | 12.889.765 | 12.609.379 | 280.386 | 33.253 | 480 | 310 | 362 | 1.193 | 791 |
| SRR11363966 | Female | no | Sedentary 4 | Healthy | Plasma | Li et al. 2020 | 19.089.860 | 5.144.366 | 4.741.002 | 403.364 | 49.296 | 426 | 516 | 303 | 1.099 | 736 |
| SRR11363967 | Female | Swimmer | Active 10 | Healthy | Plasma | Li et al. 2020 | 29.155.468 | 8.514.649 | 7.504.116 | 1.010.533 | 97.826 | 570 | 1.282 | 340 | 1.225 | 782 |
| SRR11363968 | Female | Swimmer | Active 9 | Healthy | Plasma | Li et al. 2020 | 19.739.731 | 5.560.122 | 5.006.241 | 553.881 | 46.135 | 414 | 842 | 294 | 1.092 | 754 |
| SRR11363969 | Female | Swimmer | Active 8 | Healthy | Plasma | Li et al. 2020 | 24.382.847 | 10.664.351 | 10.004.135 | 660.216 | 71.921 | 536 | 790 | 363 | 1.296 | 822 |
| SRR11363970 | Female | Swimmer | Active 7 | Healthy | Plasma | Li et al. 2020 | 28.448.347 | 13.723.589 | 12.829.709 | 893.880 | 62.432 | 587 | 760 | 377 | 1.397 | 847 |
| SRR11363971 | Female | Swimmer | Active 6 | Healthy | Plasma | Li et al. 2020 | 29.317.758 | 12.122.149 | 11.106.501 | 1.015.648 | 121.754 | 708 | 1.424 | 409 | 1.317 | 793 |
| SRR11363972 | Female | no | Sedentary 3 | Healthy | Plasma | Li et al. 2020 | 20.973.415 | 3.032.547 | 2.767.538 | 265.009 | 21.396 | 283 | 250 | 216 | 869 | 630 |
| SRR11363973 | Female | Swimmer | Active 5 | Healthy | Plasma | Li et al. 2020 | 22.113.129 | 8.891.970 | 8.211.132 | 680.838 | 57.064 | 503 | 666 | 357 | 1.201 | 781 |
| SRR11363974 | Female | Swimmer | Active 4 | Healthy | Plasma | Li et al. 2020 | 22.734.855 | 9.682.360 | 8.983.974 | 698.386 | 76.987 | 551 | 878 | 360 | 1.314 | 805 |
| SRR11363975 | Female | Swimmer | Active 3 | Healthy | Plasma | Li et al. 2020 | 21.908.930 | 15.070.142 | 13.153.194 | 1.916.948 | 107.962 | 537 | 1.492 | 330 | 1.209 | 672 |
| SRR11363976 | Female | Swimmer | Active 2 | Healthy | Plasma | Li et al. 2020 | 25.232.880 | 10.955.312 | 10.094.551 | 860.761 | 82.524 | 570 | 958 | 357 | 1.291 | 715 |
| SRR11363977 | Female | Swimmer | Active 1 | Healthy | Plasma | Li et al. 2020 | 16.552.141 | 7.801.434 | 7.232.046 | 569.388 | 44.155 | 462 | 418 | 326 | 1.204 | 770 |
| SRR11363978 | Female | no | Sedentary 15 | Healthy | Plasma | Li et al. 2020 | 14.839.099 | 1.752.536 | 1.553.728 | 198.808 | 13.192 | 221 | 228 | 175 | 767 | 595 |
| SRR11363979 | Female | no | Sedentary 14 | Healthy | Plasma | Li et al. 2020 | 17.608.864 | 12.656.961 | 10.040.941 | 2.616.020 | 193.623 | 713 | 1.838 | 401 | 1.233 | 677 |
| SRR11363980 | Female | no | Sedentary 13 | Healthy | Plasma | Li et al. 2020 | 20.914.126 | 6.552.894 | 6.260.245 | 292.649 | 24.286 | 425 | 246 | 332 | 1.130 | 761 |
| SRR11363981 | Female | no | Sedentary 12 | Healthy | Plasma | Li et al. 2020 | 21.476.762 | 9.228.210 | 8.574.123 | 654.087 | 62.937 | 607 | 760 | 395 | 1.373 | 830 |
| SRR11363982 | Female | no | Sedentary 11 | Healthy | Plasma | Li et al. 2020 | 19.411.413 | 8.231.392 | 7.787.322 | 444.070 | 40.941 | 521 | 478 | 369 | 1.288 | 837 |
| SRR11363983 | Female | no | Sedentary 2 | Healthy | Plasma | Li et al. 2020 | 11.695.923 | 11.676.363 | 11.145.105 | 531.258 | 49.776 | 636 | 422 | 430 | 1.440 | 850 |
| SRR11363984 | Female | no | Sedentary 1 | Healthy | Plasma | Li et al. 2020 | 17.510.813 | 11.106.736 | 10.709.914 | 396.822 | 49.164 | 552 | 508 | 386 | 1.324 | 826 |
| | | | | | | Total | 530.231.687 | 226.420.290 | 208.443.867 | 17.976.423 | 1.624.439 | 12.987 | 18.280 | 8.713 | 30.402 | 19.118 |
| | | | | | | | | | | | | | Unique sequences | 3.346 | 3.461 | 5.868 |

**Supplementary Table 1.** ***Sample metadata and preprocessing summary for the plasma sRNA-seq dataset used for initial small RNA candidate discovery***. The table summarizes sample metadata and preprocessing statistics for the only publicly available plasma small RNA-seq dataset suitable for candidate miRNA discovery in the context of exercise physiology. The dataset consists of young healthy female participants, including active synchronized swimmers and sedentary individuals, as originally reported by Li et al. (2020). Raw sequencing reads were processed using the nf-Sarcopipe preprocessing module, including FastQC quality assessment, adapter trimming and filtering with *fastp*, and miRNA and small RNA candidate annotation using BrumiR-core and miRDeep2. For each sample, the table reports sample identifiers, participant characteristics, tissue source (plasma), sequencing depth before and after trimming, and the number of small RNA candidates detected by each discovery method.

| Sample | Raw reads | Reads retained | Retained (%) | Lost (%) | Low-quality reads | Too short reads | Adapter-trimmed reads | Adapter | Overrepresented seq. (%) | Duplication (%) |
|---|---|---|---|---|---|---|---|---|---|---|
| SRR11363960 | 19,022,725 | 9,845,947 | 51.76 | 48.24 | 101 | 9,176,677 | 9,177,871 | PASS | 33.19 | 97.37 |
| SRR11363961 | 20,590,468 | 8,501,859 | 41.29 | 58.71 | 114 | 12,088,495 | 12,093,331 | PASS | 38.39 | 99.01 |
| SRR11363962 | 23,587,262 | 13,814,661 | 58.57 | 41.43 | 150 | 9,772,451 | 9,773,629 | PASS | 25.85 | 98.11 |
| SRR11363963 | 20,654,567 | 6,128,209 | 29.67 | 70.33 | 189 | 14,526,169 | 14,525,810 | PASS | 46.73 | 96.75 |
| SRR11363964 | 23,241,588 | 2,871,766 | 12.36 | 87.64 | 109 | 20,369,713 | 20,360,471 | PASS | 56.58 | 97.21 |
| SRR11363965 | 20,028,716 | 12,889,765 | 64.36 | 35.64 | 134 | 7,138,817 | 7,146,667 | PASS | 22.32 | 98.83 |
| SRR11363966 | 19,089,860 | 5,144,366 | 26.95 | 73.05 | 168 | 13,945,326 | 13,948,065 | PASS | 46.34 | 98.04 |
| SRR11363967 | 29,155,468 | 8,514,649 | 29.20 | 70.80 | 192 | 20,640,627 | 20,606,544 | PASS | 45.34 | 97.32 |
| SRR11363968 | 19,739,731 | 5,560,122 | 28.17 | 71.83 | 116 | 14,179,493 | 14,181,180 | PASS | 46.98 | 97.92 |
| SRR11363969 | 24,382,847 | 10,664,351 | 43.74 | 56.26 | 244 | 13,718,252 | 13,708,970 | PASS | 34.98 | 97.90 |
| SRR11363970 | 28,448,347 | 13,723,589 | 48.24 | 51.76 | 281 | 14,724,477 | 14,729,371 | PASS | 32.15 | 98.41 |
| SRR11363971 | 29,317,758 | 12,122,149 | 41.35 | 58.65 | 141 | 17,195,468 | 17,186,558 | PASS | 37.75 | 97.26 |
| SRR11363972 | 20,973,415 | 3,032,547 | 14.46 | 85.54 | 127 | 17,940,741 | 17,953,953 | PASS | 51.49 | 99.17 |
| SRR11363973 | 22,113,129 | 8,891,970 | 40.21 | 59.79 | 78 | 13,221,081 | 13,225,909 | PASS | 37.47 | 98.18 |
| SRR11363974 | 22,734,855 | 9,682,360 | 42.59 | 57.41 | 104 | 13,052,391 | 13,055,588 | PASS | 35.54 | 97.47 |
| SRR11363975 | 21,908,930 | 15,070,142 | 68.79 | 31.21 | 304 | 6,838,484 | 6,819,095 | PASS | 16.17 | 95.80 |
| SRR11363976 | 25,232,880 | 10,955,312 | 43.42 | 56.58 | 258 | 14,277,310 | 14,261,068 | PASS | 35.66 | 97.68 |
| SRR11363977 | 16,552,141 | 7,801,434 | 47.13 | 52.87 | 170 | 8,750,537 | 8,748,388 | PASS | 32.09 | 97.96 |
| SRR11363978 | 14,839,099 | 1,752,536 | 11.81 | 88.19 | 86 | 13,086,477 | 13,088,705 | PASS | 57.70 | 99.15 |
| SRR11363979 | 17,608,864 | 12,656,961 | 71.88 | 28.12 | 131 | 4,951,772 | 4,950,477 | PASS | 18.00 | 91.60 |
| SRR11363980 | 20,914,126 | 6,552,894 | 31.33 | 68.67 | 242 | 14,360,990 | 14,368,331 | PASS | 40.45 | 99.11 |
| SRR11363981 | 21,476,762 | 9,228,210 | 42.97 | 57.03 | 330 | 12,248,222 | 12,247,111 | PASS | 35.67 | 97.72 |
| SRR11363982 | 19,411,413 | 8,231,392 | 42.40 | 57.60 | 97 | 11,179,924 | 11,182,153 | PASS | 38.44 | 98.40 |
| SRR11363983 | 11,695,923 | 11,676,363 | 99.83 | 0.17 | 139 | 19,421 | 0 | PASS | 9.74 | 97.30 |
| SRR11363984 | 17,510,813 | 11,106,736 | 63.43 | 36.57 | 283 | 6,403,794 | 6,409,746 | PASS | 23.03 | 97.69 |

**Supplementary Table 2.** ***Summary of raw library quality and fastp filtering statistics across the 25 small RNA-seq libraries***. Summary of sequencing quality and preprocessing metrics obtained from FastQC and *fastp* reports for all plasma small RNA-seq libraries. For each sample, the table shows the number of raw reads, reads retained after adapter trimming, retention and loss percentages, reads discarded because of low quality or insufficient length, reads subjected to adapter trimming, FastQC adapter-content status, percentage of the most abundant overrepresented sequence, and estimated sequence duplication level. *fastp* statistics were extracted from the corresponding JSON reports generated using the same preprocessing parameters applied in the nf-Sarcopipe workflow.

| Candidate | Group | Chromosome | Start | End | Mature sequence (5'→3') | Mature length (nt) | Precursor length (nt) | BrumiR2Reference MFE (kcal/mol) | Known exact match | De novo exact |
|---|---|---|---|---|---|---|---|---|---|---|
| cluster_270 | athlete | 17 | 59137755 | 59137868 | CAGCAGAGACAAUAUUGAUAGGGU | 24 | 113 | -18.90 | 0 | 1 |
| cluster_396 | athlete | X | 50013235 | 50013347 | UACCCAUUGCAUAUCGGAGUUGU | 23 | 112 | -34.60 | 1 | 0 |
| cluster_740 | athlete | 22 | 40956683 | 40956794 | GAGCCACCAUGUCCGACCUCAA | 22 | 111 | -63.10 | 0 | 1 |
| cluster_962 | athlete | 11 | 122152203 | 122152311 | CACAAGUUCGGAUCUACGGG | 20 | 108 | -31.70 | 0 | 1 |
| cluster_986 | athlete | 17 | 58331214 | 58331324 | AGUGCUUUCUACUUUAUGGG | 20 | 110 | -39.70 | 0 | 1 |
| cluster_421 | sedentary | 14 | 101027080 | 101027192 | AACACACCUGGUUAACCUCUUUU | 23 | 112 | -36.30 | 0 | 1 |
| cluster_481 | sedentary | 14 | 101047878 | 101047990 | UUAAUAUCGGACAACCAUUGUUU | 23 | 112 | -31.00 | 0 | 1 |
| cluster_2 | shared | 3 | 49020131 | 49020240 | ACUCAACGGGAGUGAUCGUGUCAUU | 25 | 109 | -43.50 | 0 | 1 |
| cluster_42 | shared | 7 | 129770427 | 129770535 | CAGUGUGAGUUCUACCAUUGCCAAA | 25 | 108 | -40.70 | 0 | 1 |
| cluster_70 | shared | X | 151958557 | 151958671 | CUAAACGGAACCACUAGUGACUUGA | 25 | 114 | -44.50 | 0 | 1 |
| cluster_83 | shared | 14 | 101055392 | 101055504 | AGAGAGGCUGGCCGUGAUGAAUUCG | 25 | 112 | -38.50 | 0 | 1 |
| cluster_106 | shared | 1 | 40754346 | 40754452 | CUGUAAACAUCCUUGACUGGAAGCU | 25 | 106 | -54.40 | 0 | 1 |
| cluster_164 | shared | 9 | 83969750 | 83969861 | AACAACAAAAUCACUAGUCUUCCA | 24 | 111 | -36.60 | 1 | 0 |
| cluster_167 | shared | 14 | 100109644 | 100109757 | UCUCACACAGAAAUCGCACCCGUC | 24 | 113 | -50.30 | 0 | 1 |
| cluster_244 | shared | 1 | 198859037 | 198859150 | ACUCACCGACAGCGUUGAAUGUUC | 24 | 113 | -28.00 | 0 | 1 |
| cluster_374 | shared | 13 | 50048955 | 50049067 | ACGCCAAUAUUUACGUGCUGCUA | 23 | 112 | -28.10 | 0 | 1 |
| cluster_386 | shared | 11 | 57641181 | 57641293 | CAGUGCAAUGUUAAAAGGGCAUU | 23 | 112 | -47.30 | 0 | 1 |
| cluster_413 | shared | X | 134169310 | 134169417 | UACAGAUGGAUACCGUGCAAUUU | 23 | 107 | -18.90 | 0 | 1 |
| cluster_430 | shared | 9 | 124692458 | 124692571 | AACCACUGACCGUUGACUGUACC | 23 | 113 | -46.30 | 0 | 1 |
| cluster_431 | shared | 9 | 109046210 | 109046321 | UGCAACUUAGUAAUGUGCAAUAU | 23 | 111 | -23.60 | 0 | 1 |
| cluster_465 | shared | 3 | 160404733 | 160404845 | CACCAAUAUUACUGUGCUGCUUU | 23 | 112 | -36.90 | 0 | 1 |
| cluster_485 | shared | 7 | 32732980 | 32733092 | CAGUGCCUGAGGGAGUAAGAGCC | 23 | 112 | -55.80 | 0 | 1 |
| cluster_554 | shared | 8 | 134800510 | 134800622 | UGAAGUAAACAUCCACCUCCCAG | 23 | 112 | -29.30 | 0 | 1 |
| cluster_612 | shared | 2 | 135665375 | 135665486 | UCACAGUGAACCGGUCUCUUUU | 22 | 111 | -39.50 | 0 | 1 |
| cluster_650 | shared | 7 | 100093981 | 100094094 | UAUCUGCACUGUCAGCACUUUA | 22 | 113 | -48.60 | 0 | 1 |
| cluster_652 | shared | 16 | 1734966 | 1735077 | GUGCACGGCACUGGGGACACGU | 22 | 111 | -50.40 | 0 | 1 |
| cluster_819 | shared | 17 | 12081888 | 12081998 | UGCGGGGCUAGGGCUAACAGC | 21 | 110 | -41.10 | 0 | 1 |

**Supplementary Table 3.** ***Unique structurally supported BrumiR-RF core candidate miRNAs identified after BrumiR2Reference analysis.*** The table summarizes 27 unique BrumiR-RF core candidates retained after structural validation. For candidates with multiple predicted genomic loci, the locus associated with the most stable precursor, defined as the lowest minimum free energy (MFE), was retained. Reported coordinates correspond to the predicted genomic location of the precursor hairpin. The mature sequence (5′→3′) represents the predicted mature miRNA, whereas precursor length indicates the length of the predicted hairpin sequence. **Known exact match = 1** identifies candidates with an exact mature-sequence match to miRBase and/or MirGeneDB, whereas ***de novo* exact = 1** identifies structurally supported candidates lacking an exact reference match and retained as high-confidence *de novo* miRNA candidates.

| Candidate | Cluster | Length (nt) | Mature sequence (5′→3′) | Canonical seed (2–8) | log2FC | Adjusted p value | Regulation | Exact miRBase seed match | Matching miRBase family | Total 7-mers | Matching known 7-mers | Matching 7-mer(s) | Families matched across all positions |
|---|---|---|---|---|---|---|---|---|---|---|---|---|---|
| **hsa-miR-novel_A** | cluster_740 | 22 | GAGCCACCAUGUCCGACCUCAA | AGCCACC | −12.780 | $5.60 \times 10^{-14}$ | Up in sedentary | No | – | 16 | 0 | – | – |
| **hsa-miR-novel_B** | cluster_986 | 20 | AGUGCUUUCUACUUUAUGGG | GUGCUUU | −6.138 | $3.15 \times 10^{-2}$ | Up in sedentary | No | – | 14 | 2 | AGUGCUU; GCUUUCU | miR-3160-5p; miR-520f-3p |
| **hsa-miR-novel_C** | cluster_42 | 25 | CAGUGUGAGUUCUACCAUUGCCAAA | AGUGUGA | −3.359 | $8.23 \times 10^{-3}$ | Up in sedentary | No | – | 19 | 3 | AGUUCUA; CAGUGUG; CAUUGCC | miR-3168; miR-4302; miR-4317 |
| **hsa-miR-novel_D** | cluster_374 | 23 | ACGCCAAUAUUUACGUGCUGCUA | CGCCAAU | −2.339 | $3.40 \times 10^{-4}$ | Up in sedentary | No | – | 17 | 1 | CAAUAUU | miR-16-2-3p; miR-195-3p |
| **hsa-miR-novel_E** | cluster_413 | 23 | UACAGAUGGAUACCGUGCAAUUU | ACAGAUG | −2.697 | $3.55 \times 10^{-5}$ | Up in sedentary | Yes | miR-5683 | 17 | 3 | ACAGAUG; CCGUGCA; GAUGGAU | miR-1973; miR-3134; miR-5683 |
| **hsa-miR-novel_F** | cluster_465 | 23 | CACCAAUAUUACUGUGCUGCUUU | ACCAAUA | −3.810 | $3.14 \times 10^{-9}$ | Up in sedentary | No | – | 17 | 4 | AUUACUG; CAAUAUU; CUGUGCU; UACUGUG | miR-12132; miR-16-2-3p; miR-195-3p; miR-4693-5p; miR-5008-3p; miR-6737-3p; miR-7157-3p |
| **hsa-miR-novel_G** | cluster_485 | 23 | CAGUGCCUGAGGGAGUAAGAGCC | AGUGCCU | −3.221 | $7.64 \times 10^{-4}$ | Up in sedentary | Yes | miR-33b-3p; miR-515-3p; miR-519e-3p | 17 | 5 | AGUGCCU; CCUGAGG; GAGGGAG; GUGCCUG; UGAGGGA | miR-1271-3p; miR-33b-3p; miR-4510; miR-515-3p; miR-519e-3p; miR-544b; miR-550a-3-5p; miR-550a-5p; miR-6127; miR-6129; miR-6130; miR-6133; miR-6834-5p |

**Supplementary Table 4.** ***Characteristics of the final high-confidence de novo miRNA candidates identified by nf-Sarcopipe.*** Final BrumiR-derived de novo miRNA candidates retained after structural prioritization and differential expression analysis. For each candidate, the table reports the provisional miRNA identifier, original cluster, mature sequence length, mature sequence, canonical seed sequence (positions 2–8), differential expression statistics (log2 fold change, adjusted *p* value, and regulation), canonical seed match to known human miRNAs in miRBase, corresponding matched miRNA family (when applicable), and the results of the comprehensive 7-mer seed-space analysis, including the total number of possible 7-mers, the number of known matching 7-mers, the matched 7-mer sequences, and the corresponding miRNA families identified across all positions of the mature sequence.

| ID | Sex | Excercise | State | Health status | Biopsy mRNA | Raw reads (paired-end) | Reads after trimming (fastp) | Reads removed during trimming | Reference |
|---|---|---|---|---|---|---|---|---|---|
| SRR13442895_1.fastq | Female | Skater 1 - before exercise | Active | Healthy | Whole blood | 29488902 | 29306765 | 182137 | Glotov et al. 2022 |
| SRR13442895_2.fastq | Female | Skater 1 - before exercise | Active | Healthy | Whole blood | 29488902 | 29306765 | 182137 | Glotov et al. 2022 |
| SRR13442897_1.fastq | Female | Skater 2 - before exercise | Active | Healthy | Whole blood | 29895997 | 29679635 | 216362 | Glotov et al. 2022 |
| SRR13442897_2.fastq | Female | Skater 2 - before exercise | Active | Healthy | Whole blood | 29895997 | 29679635 | 216362 | Glotov et al. 2022 |
| SRR13442899_1.fastq | Female | Skater 3 - before exercise | Active | Healthy | Whole blood | 38315307 | 38102268 | 213039 | Glotov et al. 2022 |
| SRR13442899_2.fastq | Female | Skater 3 - before exercise | Active | Healthy | Whole blood | 38315307 | 38102268 | 213039 | Glotov et al. 2022 |
| SRR13442901_1.fastq | Female | Skater 4 - before exercise | Active | Healthy | Whole blood | 40888862 | 40719776 | 169086 | Glotov et al. 2022 |
| SRR13442901_2.fastq | Female | Skater 4 - before exercise | Active | Healthy | Whole blood | 40888862 | 40719776 | 169086 | Glotov et al. 2022 |
| SRR13442903_1.fastq | Female | Skater 5 - before exercise | Active | Healthy | Whole blood | 38266305 | 38077449 | 188856 | Glotov et al. 2022 |
| SRR13442903_2.fastq | Female | Skater 5 - before exercise | Active | Healthy | Whole blood | 38266305 | 38077449 | 188856 | Glotov et al. 2022 |
| SRR13442905_1.fastq | Female | Skater 6 - before exercise | Active | Healthy | Whole blood | 42600787 | 42459876 | 140911 | Glotov et al. 2022 |
| SRR13442905_2.fastq | Female | Skater 6 - before exercise | Active | Healthy | Whole blood | 42600787 | 42459876 | 140911 | Glotov et al. 2022 |
| SRR13442907_1.fastq | Female | Skater 7 - before exercise | Active | Healthy | Whole blood | 12447565 | 12323017 | 124548 | Glotov et al. 2022 |
| SRR13442907_2.fastq | Female | Skater 7 - before exercise | Active | Healthy | Whole blood | 12447565 | 12323017 | 124548 | Glotov et al. 2022 |
| SRR1424731_1.fastq | Female | no | Sedentary | Healthy | Vastus lateralis | 14977661 | 14611225 | 366436 | Lindholm et al. 2014 |
| SRR1424731_2.fastq | Female | no | Sedentary | Healthy | Vastus lateralis | 14977661 | 14611225 | 366436 | Lindholm et al. 2014 |
| SRR1424738_1.fastq | Female | no | Sedentary | Healthy | Vastus lateralis | 14174465 | 13336400 | 838065 | Lindholm et al. 2014 |
| SRR1424738_2.fastq | Female | no | Sedentary | Healthy | Vastus lateralis | 14174465 | 13336400 | 838065 | Lindholm et al. 2014 |
| SRR1424739_1.fastq | Female | no | Sedentary | Healthy | Vastus lateralis | 26183362 | 25737390 | 445972 | Lindholm et al. 2014 |
| SRR1424739_2.fastq | Female | no | Sedentary | Healthy | Vastus lateralis | 26183362 | 25737390 | 445972 | Lindholm et al. 2014 |
| SRR1424741_1.fastq | Female | no | Sedentary | Healthy | Vastus lateralis | 14288905 | 13868580 | 420325 | Lindholm et al. 2014 |
| SRR1424741_2.fastq | Female | no | Sedentary | Healthy | Vastus lateralis | 14288905 | 13868580 | 420325 | Lindholm et al. 2014 |
| SRR1424745_1.fastq | Female | no | Sedentary | Healthy | Vastus lateralis | 18265631 | 15605785 | 2659846 | Lindholm et al. 2014 |
| SRR1424745_2.fastq | Female | no | Sedentary | Healthy | Vastus lateralis | 18265631 | 15605785 | 2659846 | Lindholm et al. 2014 |
| SRR1424754_1.fastq | Female | no | Sedentary | Healthy | Vastus lateralis | 31172495 | 30154181 | 1018314 | Lindholm et al. 2014 |
| SRR1424754_2.fastq | Female | no | Sedentary | Healthy | Vastus lateralis | 31172495 | 30154181 | 1018314 | Lindholm et al. 2014 |

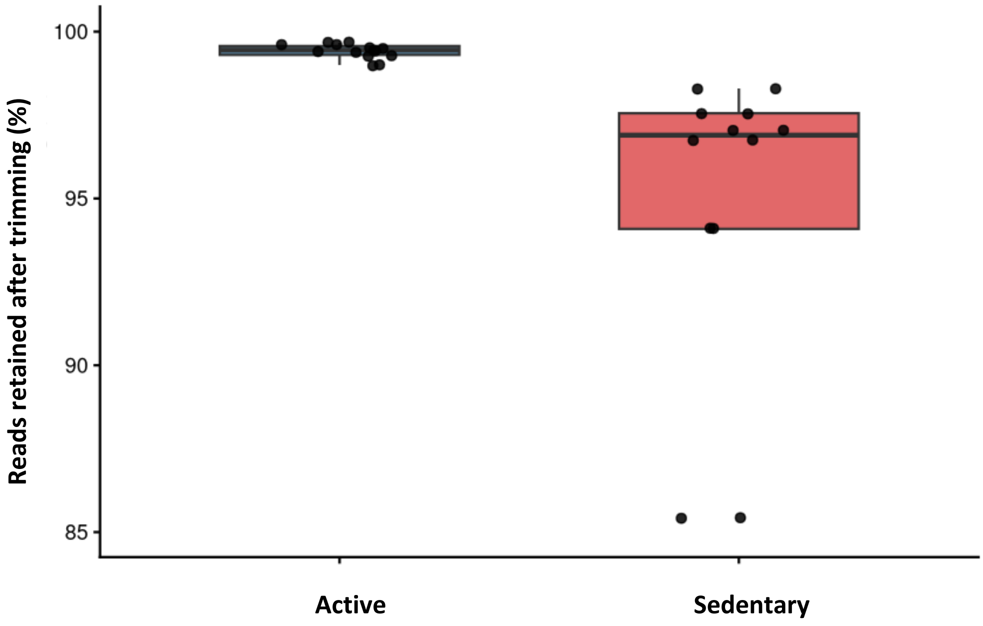

**Supplementary Table 5. *Metadata and sequencing quality metrics for the RNA-seq datasets used in the integrative analysis.*** The table summarizes sample metadata and sequencing quality metrics for RNA-seq datasets retrieved from the Gene Expression Omnibus (GEO) using GEOquery (GEPREP) and processed with the nf-core/rnaseq pipeline using the GRCh38 human reference genome. For each sample, the table reports sample identifiers (SRA accession), sex, exercise condition, activity status, health status, tissue source (whole blood or vastus lateralis muscle biopsy), and sequencing depth before and after adapter trimming using fastp for paired-end reads. The difference column indicates the number of reads removed during trimming. The datasets include RNA-seq samples from healthy young active female skaters prior to exercise and healthy sedentary female individuals, derived from previously published studies indicated in the reference column. The lower panel shows the percentage of reads retained after trimming, highlighting consistently high sequencing quality across samples.

| Gene | Functional category | Biological role | Key reference |
|---|---|---|---|
| **AKT1** | Anabolism | Central node in protein synthesis pathway | Glass 2005 |
| **CDKN1A** | Senescence | Cell cycle inhibition (p21) | Kim et al. 2025 |
| **CDKN2A** | Senescence | Cell cycle arrest (p16) | Sousa-Victor et al. 2014 |
| **COL1A1** | Fibrosis / ECM | Collagen deposition, fibrosis | Kim et al. 2025 |
| **COL3A1** | Fibrosis / ECM | Extracellular matrix remodeling | Stearns-Reider et al. 2017 |
| **COX7A1** | Mitochondrial metabolism | Electron transport chain (Complex IV) | García-Poyatos et al. 2024 |
| **FBXO32** | Atrophy / proteolysis | Ubiquitin ligase (Atrogin-1), muscle protein degradation | Gomes et al. 2001 |
| **FOXO3A** | Atrophy regulation | Transcription factor activating atrophy genes | Bogućka et al. 2025 |
| **IGF1** | Anabolism | Supports muscle growth signaling | Ruan et al. 2005 |
| **IL1B** | Inflammation | Cytokine driving chronic inflammation | Schaap et al. 2006 |
| **IL6** | Inflammation | Pro-inflammatory cytokine (inflammaging) | Visser et al. 2002 |
| **MYOD1** | Regeneration | Myogenic differentiation regulator | Raue et al. 2006 |
| **MYOG** | Regeneration | Late myogenesis marker | Raue et al. 2006 |
| **PPARGC1A** | Mitochondrial metabolism | Master regulator of mitochondrial biogenesis | Wenz et al. 2009 |
| **TNF** | Inflammation | Pro-inflammatory signaling, catabolic effects | Visser et al. 2002 |
| **TRIM63** | Atrophy / proteolysis | Ubiquitin ligase (MuRF1), sarcomere breakdown | Bodine et al. 2001 |

**Supplementary Table 6. *Functional classification of sarcopenia-associated genes***.Genes were grouped into biologically relevant categories associated with sarcopenia, including muscle atrophy, mitochondrial dysfunction, inflammation, fibrosis, regeneration, neuromuscular junction integrity, and cellular senescence. This classification is based on well-established molecular mechanisms reported in the literature, where aging skeletal muscle is characterized by increased proteolysis (e.g., FBXO32, TRIM63), chronic inflammation (e.g., IL6, TNF), and fibrosis (e.g., COL1A1, TGFB1), together with reduced mitochondrial function (e.g., PPARGC1A) and impaired regenerative capacity (e.g., PAX7, MYOD1). References supporting each functional category are provided in the corresponding column.

| ID | Condition | Function & Association (miRBase) | Function & Association (Literature - Ref) | Type of exercise | Exercise | Intervention | Expression | Biopsy | Reference |
|---|---|---|---|---|---|---|---|---|---|
| hsa-let-7b-5p | Female synchronized swimming athletes, approximately 19 years old | Predominant mature. Regulates development and cell differentiation; tumor suppressor. | Strong association with long-term training status. | Cardiorespiratory endurance | N/A (blood sample taken in the morning) | Chronic | Up | Plasma | Li et al. 2020 |
| hsa-miR-1 | Young adults, 20 app. years old | Muscle-specific miRNA essential for myogenesis and cardiac function. | Decreased after sprint. | Cardiorespiratory endurance | Two 30-second intervals of sprint cycling (SIC) | Acute | Down | Plasma | Cui et al. 2015 |
| | Young adults, 20 app. years old | | Activation of muscle-related processes. | Cardiorespiratory endurance | HIIE: high-intensity interval running, followed by vigorous continuous exercise after 3 days. | Acute | Up | Plasma | Cui et al. 2016 |
| | Young adults, 20 app. years old | | ↑ in endurance athletes (long-term), ↓ immediately after maximal exercise (acute). | Cardiorespiratory endurance | Single maximal aerobic exercise session | Acute<br>Chronic | Down<br>UP | Whole blood | Denham et al. 2016 |
| | Young adults aged 26 to 37 years | | Increase only after eccentric exercise | Cardiorespiratory endurance | Two walking protocols (6-week gap): downhill eccentric, uphill concentric | Acute | Up | Plasma | Banzet et al. 2013 |
| | Young adults, 22 app. years old | | Responds to increasing intensity of exercise | Cardiorespiratory endurance | Two treadmill protocols: variable intensity and duration | Acute | Up | Plasma | Ramos et al. 2018 |
| | Male marathon runners, average age 32 years | | Significantly increased after a half-marathon, indicating muscle stress/adaptation. | Cardiorespiratory endurance | Half marathon | Acute | Up | Plasma | Gomes et al. 2014 |
| | Healthy women and men aged 51 to 56 years | | Rise after marathon running, reflecting normal muscle stress. | Cardiorespiratory endurance | Marathon | Acute | Up | Plasma | Pil-Gi Min et al. 2015 |
| | 21 healthy male marathon runners | | Muscle-enriched, upregulated after marathon. | Cardiorespiratory endurance | Marathon | Acute | Up | Plasma | Baggish et al. 2014 |
| | Male marathon runners, average age app. 42 years. | | Muscle- and heart-specific → significantly increased after the marathon. | Cardiorespiratory endurance | Marathon | Acute | Up | Plasma | Mooren et al. 2014 |
| | Adult men, approximately 33 years old | | In this short-term SIT intervention, their circulating levels decreased. | Cardiorespiratory endurance | 8 weeks of single sprint interval training (3 times per week) | Acute | Down | Whole blood | Denham et al. 2017 |
| | Elite endurance athletes, men, approximately 60 years old | | Increased significantly. | Cardiorespiratory endurance | 11 weeks of endurance and marathon training | Acute | Up | Plasma | Clauss et al. 2016 |
| | Healthy sedentary young adults | | Lower expression after exercise. | Cardiorespiratory endurance | Six 45-minute cycling sessions per week (6) | Chronic | Down | Muscle (vastus lateralis) | Keller et al. 2011 |
| hsa-miR-1-3p | Young men, average age 24.6 ± 4.0 years. | Predominant mature strand of miR-1; regulates muscle differentiation. | Exercise-induced release of exosomes enriched in specific miRNAs. | Cardiorespiratory endurance | One session of high-intensity interval cycling exercise | Acute | Down | Plasma | D'Souza et al. 2018 |
| | Young men, average age 24.6 ± 4.0 years. | | Exercise-induced release of exosomes enriched in specific miRNAs. | Cardiorespiratory endurance | One session of high-intensity interval cycling exercise | Acute | Up | Exosomes | D'Souza et al. 2018 |
| | Adults app. 31 years old | | Normal response of muscle-specific miRNAs to acute muscle damage. | Cardiorespiratory endurance | Two downhill treadmill sessions (2-week gap) causing muscle damage | Acute | Up | Plasma | Chalchat et al. 2022 |
| | Young adults, 20 app. years old | | Overexpressed immediately, returned to baseline at 24 h. | Cardiorespiratory endurance | 8 km run | Acute | Up | Serum | Yin et al. 2020 |
| | Elite endurance athletes, men and women, approximately 45 years old | | Strongly overexpressed, reflects acute muscle damage. | Cardiorespiratory endurance | 24-hour World Running Championship (exhaustive running causing muscle damage) | Acute | Up | Plasma | Chalchat et al. 2021 |
| | Elite endurance athletes, men and women, approximately 45 years old | | Correlate with muscle function loss. | Cardiorespiratory endurance | 24-hour World Running Championship (exhaustive running causing muscle damage) | Acute | Up | Plasma | Chalchat et al. 2021 |
| hsa-miR-16 | Young adults, 20 app. years old | Ubiquitously expressed tumor-suppressor group. | Systemic somatic stress response. | Cardiorespiratory endurance | Two 30-second intervals of sprint cycling (SIC) | Acute | Down | Plasma | Cui et al. 2015 |
| hsa-miR-101 | Healthy sedentary young adults | Tumor suppressor; regulates proliferation and apoptosis. | Lower expression after exercise. | Cardiorespiratory endurance | Six 45-minute cycling sessions per week (6) | Chronic | Down | Muscle (vastus lateralis) | Keller et al. 2011 |
| hsa-miR-103 | Young healthy men | Regulates insulin receptor signaling and glucose uptake. | Metabolic adaptation and insulin sensitivity. | Cardiorespiratory endurance | 12 weeks of endurance training (5 times per week) | Chronic | Up | Plasma | Nielsen et al. 2014 |
| hsa-miR-107 | Young healthy men | Regulates energy metabolism and lipid/glucose homeostasis. | Metabolic adaptation and insulin sensitivity. | Cardiorespiratory endurance | 12 weeks of endurance training (5 times per week) | Chronic | Up | Plasma | Nielsen et al. 2014 |
| hsa-miR-122 | Young adults, 20 app. years old | Regulates energy and lipid metabolism (liver-specific). | Acute metabolic regulation related to liver and energy metabolism. | Cardiorespiratory endurance | Two 30-second intervals of sprint cycling (SIC) | Acute | Down | Plasma | Cui et al. 2015 |
| hsa-miR-125a | Healthy sedentary young adults | Regulates cell differentiation and proliferation. | Higher expression after exercise. | Cardiorespiratory endurance | Six 45-minute cycling sessions per week (6) | Chronic | Up | Muscle (vastus lateralis) | Keller et al. 2011 |
| hsa-miR-133 | Male marathon runners, average age 32 years | Muscle-specific; controls myogenesis and muscle repair. | Combined endothelial and muscle damage. | Cardiorespiratory endurance | Marathon | Acute | Up | Plasma | Uhlemann et al. 2012 |
| | Trained men and women, average age 37 years | | Muscle damage without endothelial injury. | Strength exercise | Acute resistance exercise session (weights) | Acute | Up | Plasma | Uhlemann et al. 2012 |
| hsa-miR-133a | Young adults aged 26 to 37 years | Regulates muscle growth and differentiation. | Increase only after eccentric exercise. | Cardiorespiratory endurance | Two walking protocols (6-week gap): downhill eccentric, uphill concentric | Acute | Up | Plasma | Banzet et al. 2013 |
| | Young adults, 20 app. years old | | The authors interpret this downregulation as an acute physiological adaptation to anaerobic stress. | Cardiorespiratory endurance | Two 30-second intervals of sprint cycling (SIC) | Acute | Down | Plasma | Cui et al. 2015 |
| | Young adults, 20 app. years old | | Activation of muscle-related processes. | Cardiorespiratory endurance | HIIE: high-intensity interval running, followed by vigorous continuous exercise after 3 days. | Acute | Up | Plasma | Cui et al. 2016 |
| | Young adults, 20 app. years old | | ↓ immediately after maximal exercise. | Cardiorespiratory endurance | Single maximal aerobic exercise session | Acute | Down | Whole blood | Denham et al. 2016 |
| | Young adults, 22 app. years old | | Responds to increasing duration of exercise. | Cardiorespiratory endurance | Two treadmill protocols: variable intensity and duration | Acute | Up | Plasma | Ramos et al. 2018 |
| | Male marathon runners, average age 32 years | | Significantly increased after a half-marathon, indicating muscle stress/adaptation. | Cardiorespiratory endurance | Half marathon | Acute | Up | Plasma | Gomes et al. 2014 |
| | Healthy women and men aged 51 to 56 years | | Rise after marathon running, reflecting normal muscle stress. | Cardiorespiratory endurance | Marathon | Acute | Up | Plasma | Pil-Gi Min et al. 2015 |
| | Male marathon runners, average age 51 years | | Muscle-enriched, upregulated. | Cardiorespiratory endurance | Marathon | Acute | Up | Plasma | Baggish et al. 2014 |
| | Male marathon runners, average age app. 42 years. | | Muscle- and heart-specific → significantly increased after the marathon. | Cardiorespiratory endurance | Marathon | Acute | Up | Plasma | Mooren et al. 2014 |
| | Adult men, approximately 33 years old | | In this short-term SIT intervention, their circulating levels decreased. | Cardiorespiratory endurance | 7 weeks of single sprint interval training (3 times per week) | Acute | Down | Whole blood | Denham et al. 2017 |
| | Male athlete app. 40 years old | | Increased significantly. | Cardiorespiratory endurance | 10 weeks of endurance and marathon training | Acute | Up | Plasma | Clauss et al. 2016 |
| | Elite endurance athletes, men, approximately 60 years old | | Increased significantly. | Cardiorespiratory endurance | 12 weeks of endurance and marathon training | Acute | Up | Plasma | Clauss et al. 2016 |
| | Young men, average age 19 years. | | Reflect muscle adaptation and favorable hormonal balance. | Strength exercise | Muscle hypertrophy (MH) | Acute | Up | Plasma | Cui et al. 2017 |
| | Young men, average age 19 years. | | Reflect muscle adaptation and favorable hormonal balance. | Strength exercise | Strength endurance (SE) | | Up | | |
| | Young men, average age 19 years. | | Reflect muscle adaptation and favorable hormonal balance. | Strength exercise | Maximum strength (MS) | | Up | | |
| | Trained men, average age 24 years. | | Transient and limited. | Strength exercise | Acute resistance exercise session | Acute | Up | Plasma | D'Souza et al. 2019 |
| | Healthy sedentary young adults | | Lower expression after exercise. | Cardiorespiratory endurance | Six 45-minute cycling sessions per week (6) | Chronic | Down | Muscle (vastus lateralis) | Keller et al. 2011 |
| hsa-miR-133a-3p | Adults app. 31 years old | Predominant mature strand of miR-133a; regulates muscle remodeling. | Normal response of muscle-specific miRNAs to acute muscle damage. | Cardiorespiratory endurance | Two downhill treadmill sessions (2-week gap) causing muscle damage | Acute | Up | Plasma | Chalchat et al. 2022 |
| | Young adults, 20 app. years old | | Overexpressed immediately and remained elevated at 24 h. | Cardiorespiratory endurance | 8 km run | Acute | Up | Serum | Yin et al. 2020 |
| | Elite endurance athletes, men and women, approximately 45 years old | | Strongly overexpressed. | Cardiorespiratory endurance | 24-hour World Running Championship (exhaustive running causing muscle damage) | Acute | Up | Plasma | Chalchat et al. 2021 |
| | Elite endurance athletes, men and women, approximately 45 years old | | Correlate with muscle function loss. | Cardiorespiratory endurance | 24-hour World Running Championship (exhaustive running causing muscle damage) | Acute | Up | Plasma | Chalchat et al. 2021 |
| hsa-miR-133b | Young adults, 20 app. years old | Regulates muscle and neuronal function; therapeutic potential. | Activation of muscle-related processes. | Cardiorespiratory endurance | HIIE: high-intensity interval running, followed by vigorous continuous exercise after 3 days. | Acute | Up | Plasma | Cui et al. 2016 |
| | Elite endurance athletes, men and women, approximately 45 years old | | Correlate with muscle function loss. | Cardiorespiratory endurance | 24-hour World Running Championship (exhaustive running causing muscle damage) | Acute | Up | Plasma | Chalchat et al. 2021 |
| | Young men, average age 19 years. | | Reflect muscle adaptation and favorable hormonal balance. | Strength exercise | Muscle hypertrophy (MH) | Acute | Up | Plasma | Cui et al. 2017 |
| | Young men, average age 19 years. | | Reflect muscle adaptation and favorable hormonal balance. | Strength exercise | Strength endurance (SE) | Acute | Up | Plasma | Cui et al. 2017 |
| | Adult men, approximately 33 years old | | In this short-term SIT intervention, their circulating levels decreased. | Cardiorespiratory endurance | 7 weeks of single sprint interval training (3 times per week) | Acute | Down | Whole blood | Denham et al. 2017 |
| | Young men, average age 19 years. | | Reflect muscle adaptation and favorable hormonal balance. | Strength exercise | Maximum strength (MS) | Acute | Up | Plasma | Cui et al. 2017 |
| | Young adults, 20 app. years old | | Decreased after sprint and correlated with peak power. | Cardiorespiratory endurance | Two 30-second intervals of sprint cycling (SIC) | Acute | Down | Plasma | Cui et al. 2015 |
| | Young adults aged 26 to 37 years | | Increase only after eccentric exercise | Cardiorespiratory endurance | Two walking protocols (6-week gap): downhill eccentric, uphill concentric | Acute | Up | Plasma | Banzet et al. 2013 |
| | Young adults, 20 app. years old | | Involved in muscle adaptation and remodeling. | Cardiorespiratory endurance | Two exercise protocols (8-week gap): acute and maximal aerobic sessions | Acute | Up | Exosomes | Maggio et al. 2023 |
| | Young adults, 20 app. years old | | Overexpressed immediately, returned to baseline at 24 h. | Cardiorespiratory endurance | 8 km run | Acute | Up | Serum | Yin et al. 2020 |
| | Adult men, approximately 33 years old | | Decreased after 6 weeks of sprint interval training. | Cardiorespiratory endurance | 6 weeks of single sprint interval training (3 times per week) | Acute | Down | Whole blood | Denham et al. 2018 |
| hsa-miR-144 | Healthy sedentary young adults | Regulates erythropoiesis and cell proliferation. | Lower expression after exercise. | Cardiorespiratory endurance | Six 45-minute cycling sessions per week (6) | Chronic | Down | Muscle (vastus lateralis) | Keller et al. 2011 |
| hsa-miR-146a | Young adults, 22 app. years old | Plays a key role in chondrogenesis, immune regulation, and anti-inflammatory responses. | Increases after exercise, independent of intensity/duration. | Cardiorespiratory endurance | Two treadmill protocols: variable intensity and duration | Acute | Up | Plasma | Ramos et al. 2018 |
| | Young adults, 20 app. years old | | Key regulator of inflammation. | Cardiorespiratory endurance | Two exercise protocols (8-week gap): acute and maximal aerobic sessions | Acute | Up | Exosomes | Maggio et al. 2023 |
| | | | Involved in inflammatory and immune regulation. | Strength exercise | Single flywheel training session | Acute | Up | Exosomes | Annibalini et al. 2019 |
| hsa-miR-149 | Trained men, average age 24 years. | It is involved in cell cycle and extracellular matrix regulation. | Transient and limited. | Strength exercise | Acute resistance exercise session | Acute | Up | Plasma | D'Souza et al. 2019 |
| hsa-miR-15b | Healthy sedentary young adults | Regulates cell cycle and apoptosis. | Lower expression after exercise. | Cardiorespiratory endurance | Six 45-minute cycling sessions per week (6) | Chronic | Down | Muscle (vastus lateralis) | Keller et al. 2011 |
| hsa-miR-181a | Young adults, average age 21 years | Regulates differentiation and immune response. | Upregulated 3 h after acute exercise. | Cardiorespiratory endurance | 60-minute cycling sessions at 70% VO2peak for 7 days + 4 high-intensity sessions | Acute | Up | Muscle (vastus lateralis) | Russell et al. 2013 |
| | Young men, average age 19 years. | | Reflects immune/inflammatory activation | Strength exercise | Muscle hypertrophy (MH) | Acute | Up | Plasma | Cui et al. 2017 |
| | Young men, average age 19 years. | | | Strength exercise | Strength endurance (SE) | | | | |
| hsa-miR-183 | Healthy sedentary young adults | Involved in sensory organ development and proliferation. | Higher expression after exercise. | Cardiorespiratory endurance | Six 45-minute cycling sessions per week (6) | Chronic | Up | Muscle (vastus lateralis) | Keller et al. 2011 |
| hsa-miR-189 | Healthy sedentary young adults | Regulates gene expression in development. | Higher expression after exercise. | Cardiorespiratory endurance | Six 45-minute cycling sessions per week (6) | Chronic | Up | Muscle (vastus lateralis) | Keller et al. 2011 |
| hsa-miR-199a-5p | Young adults, 22 app. years old | Acts mainly as a cell-cycle and apoptosis regulator; functions as a tumor suppressor and is linked to stress and damage responses. | Shows opposite regulation between exhaustive and non-exhaustive exercise. | Cardiorespiratory endurance | Single exhaustive endurance exercise (including 4 phases) | Chronic | Down | Plasma | Håkansson et al. 2018 |
| hsa-miR-194-5p | Elderly males aged 65–80 years | Involved is controlling proliferation and inflammatory signaling. | Enhancing mitochondrial function, energy metabolism, and metabolic efficiency. | Cardiorespiratory endurance | Long-Term endurance training. | Chronic | Up | Muscle | Zhi et al. 2025 |
| hsa-miR-206 | Young adults, 20 app. years old | Muscle-specific; promotes myogenic differentiation and regeneration. | Activation of muscle-related processes. | Cardiorespiratory endurance | HIIE: high-intensity interval running, followed by vigorous continuous exercise after 3 days. | Acute | Up | Plasma | Cui et al. 2016 |
| | Adults app. 31 years old | | Normal response of muscle-specific miRNAs to acute muscle damage. | Cardiorespiratory endurance | Two downhill treadmill sessions (2-week gap) causing muscle damage | Acute | Up | Plasma | Chalchat et al. 2022 |
| | Young adults, 20 app. years old | | Muscle-specific; strongly responsive to exercise. | Cardiorespiratory endurance | Two exercise protocols (8-week gap): acute and maximal aerobic sessions | Acute | Up | Exosomes | Maggio et al. 2023 |
| | Young adults, 20 app. years old | | Decreased at 24 h; may indicate post-exercise adaptation or delayed regulation. | Cardiorespiratory endurance | 8 km run | Acute | Up | Serum | Yin et al. 2020 |
| | Male marathon runners, average age 32 years | | Significantly increased after a half-marathon, indicating muscle stress/adaptation. | Cardiorespiratory endurance | Half marathon | Acute | Up | Plasma | Gomes et al. 2014 |
| | Healthy women and men aged 51 to 56 years | | Rise after marathon running, reflecting normal muscle stress. | Cardiorespiratory endurance | Marathon | Acute | Up | Plasma | Pil-Gi Min et al. 2015 |
| | Male marathon runners, average age app. 42 years. | | Muscle- and heart-specific → significantly increased after the marathon. | Cardiorespiratory endurance | Marathon | Acute | Up | Plasma | Mooren et al. 2014 |
| | Obese women and men post-gastric surgery, aged 36 to 48 years | | Exercise adaptation. | Cardiorespiratory endurance | 6 months of progressive endurance training (3-5 times per week) | Chronic | Up | Plasma | Nunes et al. 2017 |
| | Female synchronized swimming athletes, approximately 19 years old | | Muscle-related | Cardiorespiratory endurance | N/A (blood sample taken in the morning) | Chronic | Up | Plasma | Li et al. 2020 |
| | Trained men, average age 24 years. | | Linked to skeletal muscle remodeling, regeneration, and myogenic signaling. | Strength exercise | Single flywheel training session | Acute | Up | Exosomes | Annibalini et al. 2019 |
| | Young men, average age 19 years. | | Linked to muscle remodeling. | Strength exercise | Muscle hypertrophy (MH) | Acute | Up | Plasma | Cui et al. 2017 |
| | Young men, average age 19 years. | | Linked to muscle remodeling. | Strength exercise | Maximum strength (MS) | | Up | | |
| | Young men, average age 19 years. | | Linked to muscle remodeling. | Strength exercise | Strength endurance (SE) | | Up | | |
| hsa-miR-208a-3p | Elite endurance athletes, men and women, approximately 45 years old | Regulate cardiac muscle identity and adaptation. | Strongly overexpressed, cardiac-specific relevance. | Cardiorespiratory endurance | 24-hour World Running Championship (exhaustive running causing muscle damage) | Acute | Up | Plasma | Chalchat et al. 2021 |
| hsa-miR-208b | Male marathon runners, average age app. 42 years. | Regulates heart and muscle myosin, affects hypertrophy and fiber type, and can indicate heart disease. | Muscle- and heart-specific → significantly increased after the marathon. | Cardiorespiratory endurance | Marathon | Acute | Up | Plasma | Mooren et al. 2014 |
| | Young men, average age 19 years. | | Reflects load-specific muscle stress. | Strength exercise | Strength endurance (SE) | Acute | Up | Plasma | Cui et al. 2017 |
| hsa-miR-208b-3p | Elite endurance athletes, men and women, approximately 45 years old | Regulate cardiac muscle identity and adaptation. | Strongly overexpressed, cardiac-specific relevance. | Cardiorespiratory endurance | 24-hour World Running Championship (exhaustive running causing muscle damage) | Acute | Up | Plasma | Chalchat et al. 2021 |
| hsa-miR-221 | Young men, average age 19 years. | Regulates cell survival, apoptosis, and autophagy and acts as an oncomiR by activating PI3K/Akt/mTOR pathway | Intensity-dependent inflammatory/remodeling response | Strength exercise | Muscle hypertrophy (MH) | Acute | Up | Plasma | Cui et al. 2017 |
| hsa-miR-222 | Young adults, 22 app. years old | Regulates adipogenesis and vascular smooth muscle cell proliferation. | Responds to increasing duration of exercise. | Cardiorespiratory endurance | Two treadmill protocols: variable intensity and duration | Acute | Up | Plasma | Ramos et al. 2018 |
| hsa-miR-21 | Young healthy men | Involved in inflammation, wound healing, and tissue remodeling. | Regulator of inflammation and tissue remodeling. | Cardiorespiratory endurance | 12 weeks of endurance training (5 times per week) | Chronic | Down | Plasma | Nielsen et al. 2014 |
| | Young men, average age 19 years. | | Associated with inflammation and tissue remodeling | Strength exercise | Muscle hypertrophy (MH) | Acute | Up | Plasma | Cui et al. 2017 |
| hsa-miR-21-5p | Female synchronized swimming athletes, approximately 19 years old | Promotes cell survival, proliferation. | Strong association with long-term training status. | Cardiorespiratory endurance | N/A (blood sample taken in the morning) | Chronic | Up | Plasma | Li et al. 2020 |
| | Elderly males aged 65–80 years | | Enhance protein synthesis and reduce muscle cell apoptosis through metabolic and growth signaling pathways. | Strength exercise | Long-Term resistance exercise. | Chronic | Up | Muscle | Zhi et al. 2025 |
| hsa-miR-23a | Young adults, average age 21 years | Regulates muscle metabolism and cell differentiation. | Downregulated 3 h after acute exercise. | Cardiorespiratory endurance | 60-minute cycling sessions at 70% VO2peak for 7 days + 4 high-intensity sessions | Acute | Down | Muscle (vastus lateralis) | Russell et al. 2013 |
| hsa-miR-24 | Young adults, 22 app. years old | Influences cell proliferation, differentiation, and cancer-related pathways. | Increases after exercise, independent of intensity/duration. | Cardiorespiratory endurance | Two treadmill protocols: variable intensity and duration | Acute | Up | Plasma | Ramos et al. 2018 |
| hsa-miR-26a | Men aged 18 to 30 years | Controls proliferation and differentiation. | In low responders suggests impaired activation of muscle growth and remodeling pathways. | Strength exercise | Acute resistance exercise session (weights) | Chronic | Down | Muscle (vastus lateralis) | Davidsen et al. 2011 |
| | Elite endurance athletes, men, approximately 60 years old | | Increased significantly. | Cardiorespiratory endurance | 12 weeks of endurance and marathon training | Acute | Down | Plasma | Clauss et al. 2016 |
| hsa-miR-26b | Healthy sedentary young adults | Controls proliferation and differentiation. | Lower expression after exercise. | Cardiorespiratory endurance | Six 45-minute cycling sessions per week (6) | Chronic | Down | Muscle (vastus lateralis) | Keller et al. 2011 |
| hsa-miR-28 | Healthy sedentary young adults | Regulates proliferation and apoptosis. | Lower expression after exercise. | Cardiorespiratory endurance | Six 45-minute cycling sessions per week (6) | Chronic | Down | Muscle (vastus lateralis) | Keller et al. 2011 |
| hsa-miR-29a | Men aged 18 to 30 years | Regulates fibrosis, extracellular matrix, and apoptosis. | In low responders suggests impaired activation of muscle growth and remodeling pathways. | Strength exercise | Acute resistance exercise session (weights) | Chronic | Down | Muscle (vastus lateralis) | Davidsen et al. 2011 |

| | | | | | | | | | |
|---|---|---|---|---|---|---|---|---|---|
| hsa-miR-29a-3p | Young adults, 22 app. years old | Regulates extracellular matrix remodeling by targeting collagen and fibrosis-related genes. | ↑ after non-exhaustive exercise. | Cardiorespiratory endurance | Single exhaustive endurance exercise (including 4 phases) | Chronic | Down | Plasma | Håkansson et al. 2018 |
| hsa-miR-29b | Healthy sedentary young adults | Regulates fibrosis and cell differentiation. | Lower expression after exercise. | Cardiorespiratory endurance | Six 45-minute cycling sessions per week (6) | Chronic | Down | Muscle (vastus lateralis) | Keller et al. 2011 |
| | Young adults, average age 23 years | | Short-term training effects. | Cardiorespiratory endurance | 60-minute cycling sessions at 70% VO2peak for 7 days + 4 high-intensity sessions | Acute | Up | Muscle (vastus lateralis) | Russell et al. 2013 |
| hsa-miR-31 | Young adults, average age 23 years | Controls myogenesis and proliferation. | Short-term training effects. | Cardiorespiratory endurance | 60-minute cycling sessions at 70% VO2peak for 7 days + 4 high-intensity sessions | Acute | Down | Muscle (vastus lateralis) | Russell et al. 2013 |
| hsa-miR-34a-5p | Elderly males aged 65–80 years | Controls cell cycle arrest, apoptosis, and cellular senescence. | Enhance protein synthesis and reduce muscle cell apoptosis through metabolic and growth signaling pathways. | Strength exercise | Long-Term resistance exercise. | Chronic | Up | Muscle | Zhi et al. 2025 |
| | Elderly males aged 65–80 years | | Enhancing mitochondrial function, energy metabolism, and metabolic efficiency. | Cardiorespiratory endurance | Long-Term endurance training. | Chronic | Up | Muscle | Zhi et al. 2025 |
| hsa-miR-338 | Healthy sedentary young adults | Involved in neuronal differentiation. | Lower expression after exercise. | Cardiorespiratory endurance | Six 45-minute cycling sessions per week (6) | Chronic | Down | Muscle (vastus lateralis) | Keller et al. 2011 |
| hsa-miR-378 | Men aged 18 to 30 years | Regulates metabolism and myogenesis. | In low responders suggests impaired activation of muscle growth and remodeling pathways. | Strength exercise | Acute resistance exercise session (weights) | Chronic | Down | Muscle (vastus lateralis) | Davidsen et al. 2011 |
| hsa-miR-423-5p | Female synchronized swimming athletes, approximately 19 years old | Involved in the regulation of cell cycle, metabolism, and stress responses. | Strong association with long-term training status. | Cardiorespiratory endurance | N/A (blood sample taken in the morning) | Chronic | Up | Plasma | Li et al. 2020 |
| hsa-miR-451 | Men aged 18 to 30 years | Regulates erythropoiesis and stress response. | Reflects a compensatory response associated with limited hypertrophic adaptation. | Strength exercise | Acute resistance exercise session (weights) | Chronic | Up | Muscle (vastus lateralis) | Davidsen et al. 2011 |
| | Healthy sedentary young adults | | Lower expression after exercise. | Cardiorespiratory endurance | Six 45-minute cycling sessions per week (6) | Chronic | Down | Muscle (vastus lateralis) | Keller et al. 2011 |
| hsa-miR-451a | Female synchronized swimming athletes, approximately 19 years old | Upregulated in disease, notably active tuberculosis and glioblastoma. | Muscle-related | Cardiorespiratory endurance | N/A (blood sample taken in the morning) | Chronic | Up | Plasma | Li et al. 2020 |
| hsa-miR-455 | Healthy sedentary young adults | Regulates cartilage and muscle differentiation. | Lower expression after exercise. | Cardiorespiratory endurance | Six 45-minute cycling sessions per week (6) | Chronic | Down | Muscle (vastus lateralis) | Keller et al. 2011 |
| hsa-miR-486 | Young adults, 22 app. years old | Associated with the regulation of genes involved in cell growth and survival. | In this short-term SIT intervention, their circulating levels decreased. | Cardiorespiratory endurance | 8 weeks of single sprint interval training (3 times per week) | Acute | Down | Whole blood | Denham et al. 2017 |
| | Young adults, 20 app. years old | | ↑ in endurance athletes (long-term), ↓ immediately after maximal exercise. | Cardiorespiratory endurance | Single maximal aerobic exercise session | Chronic | Down | Whole blood | Denham et al. 2016 |
| | | | | | | Acute | UP | | |
| hsa-miR-494 | | Oncogenic miRNA that lowers cholesterol, promotes apoptosis, and regulates ATF3/JUN. | Linked to aerobic adaptation. | | | Chronic | UP | | |
| hsa-miR-495a-3p | Young adults, 22 app. years old | Involved in myogenesis and cell differentiation. | Associated with poor muscle recovery. | Cardiorespiratory endurance | Single exhaustive endurance exercise (including 4 phases) | Chronic | Down | Plasma | Håkansson et al. 2018 |
| hsa-miR-486 | Female synchronized swimming athletes, approximately 19 years old | Involved in muscle development, regeneration, and metabolism. | Muscle-related | Cardiorespiratory endurance | N/A (blood sample taken in the morning) | Chronic | Up | Plasma | Li et al. 2020 |
| hsa-miR-499 | Male marathon runners, average age app. 42 years | Muscle-specific, mainly expressed in cardiac and skeletal muscle. | Muscle- and heart-specific → significantly increased after the marathon. | Cardiorespiratory endurance | Marathon | Acute | Up | Plasma | Mooren et al. 2013 |
| miR-499a-5p | Elite endurance athletes, men and women, approximately 45 years old | Regulates muscle fiber identity (slow/oxidative) | Strongly overexpressed; cardiac-specific relevance. | Cardiorespiratory endurance | 24-hour World Running Championship (exhaustive running causing muscle damage) | Acute | Up | Plasma | Chalchat et al. 2021 |
| hsa-miR-499-5p | Healthy women and men aged 51 to 56 years | Muscle-specific, mainly expressed in cardiac and skeletal muscle. | Specifically increases with statin-related muscle injury. | Cardiorespiratory endurance | Marathon | Acute | Up | Plasma | Pål-Ki Min et al. 2015 |
| | 21 healthy male marathon runners | | Muscle-enriched, upregulated. | | | Acute | Up | Plasma | Baggish et al. 2014 |
| hsa-miR-532 | Young men, average age 19 years | Involved in cancer regulation, often acting as a tumor suppressor by limiting cell proliferation and invasion. | Associated with an anti-inflammatory/anabolic environment. Negatively correlated with IGF-1 and positively with IL-10. | Strength exercise | Muscle hypertrophy (MH) | Acute | Up | Plasma | Cui et al. 2017 |
| hsa-miR-574-5p | Elderly males aged 65–80 years | Regulation of inflammation, cell proliferation, apoptosis, and metabolic signaling. | Enhance protein synthesis and reduce muscle cell apoptosis through metabolic and growth signaling pathways. | Strength exercise | Long-Term resistance exercise. | Chronic | Up | Muscle | Zhi et al. 2025 |
| hsa-miR-575 | Healthy sedentary young adults | Regulates proliferation and apoptosis. | Higher expression after exercise. | Cardiorespiratory endurance | Six 45-minute cycling sessions per week (6) | Chronic | Up | Muscle (vastus lateralis) | Keller et al. 2011 |
| hsa-miR-589 | Healthy sedentary young adults | Regulates proliferation and migration. | Lower expression after exercise. | Cardiorespiratory endurance | Six 45-minute cycling sessions per week (6) | Chronic | Down | Muscle (vastus lateralis) | Keller et al. 2011 |
| hsa-miR-636 | Healthy sedentary young adults | Involved in cell proliferation and apoptosis. | Higher expression after exercise. | Cardiorespiratory endurance | Six 45-minute cycling sessions per week (6) | Chronic | Up | Muscle (vastus lateralis) | Keller et al. 2011 |
| hsa-miR-637 | Healthy sedentary young adults | Regulates proliferation and differentiation. | Higher expression after exercise. | Cardiorespiratory endurance | Six 45-minute cycling sessions per week (6) | Chronic | Up | Muscle (vastus lateralis) | Keller et al. 2011 |
| hsa-miR-9 | Young adults, average age 23 years | Regulates neuronal differentiation and development. | Downregulated 3 h after acute exercise. | Cardiorespiratory endurance | 60-minute cycling sessions at 70% VO2peak for 7 days + 4 high-intensity sessions | Acute | Down | Muscle (vastus lateralis) | Russell et al. 2013 |
| hsa-miR-92 | Healthy sedentary young adults | Regulates angiogenesis and proliferation. | Lower expression after exercise. | Cardiorespiratory endurance | Six 45-minute cycling sessions per week (6) | Chronic | Down | Muscle (vastus lateralis) | Keller et al. 2011 |
| hsa-miR-98 | Healthy sedentary young adults | Controls proliferation and apoptosis. | Lower expression after exercise. | Cardiorespiratory endurance | Six 45-minute cycling sessions per week (6) | Chronic | Down | Muscle (vastus lateralis) | Keller et al. 2011 |
| hsa-miR-23b | Young adults, average age 23 years | Regulates muscle metabolism and differentiation. | Downregulated 3 h after acute exercise. | Cardiorespiratory endurance | 60-minute cycling sessions at 70% VO2peak for 7 days + 4 high-intensity sessions | Acute | Down | Muscle (vastus lateralis) | Russell et al. 2013 |
| hsa-miR-422* | Healthy sedentary young adults | Minor strand; regulates proliferation and differentiation. | Higher expression after exercise. | Cardiorespiratory endurance | Six 45-minute cycling sessions per week (6) | Chronic | Up | Muscle (vastus lateralis) | Keller et al. 2011 |
| hsa-miR-6831-5p | Elderly males aged 65–80 years | Expressed in circulation and predicted to regulate genes involved in metabolic and stress-response pathways. | Enhancing mitochondrial function, energy metabolism, and metabolic efficiency. | Cardiorespiratory endurance | Long-Term endurance training. | Chronic | Up | Muscle | Zhi et al. 2025 |

**Supplementary Table 7 (page 1 & 2).** ***Curated list of microRNAs associated with exercise responses in humans***. The table summarizes microRNAs (miRNAs) previously reported to be associated with exercise responses in young and older adults, including both cardiorespiratory endurance and strength exercise paradigms. For each miRNA, the table reports the study population or condition, functional annotations from miRBase, functional associations described in the literature, exercise modality, intervention type (acute or chronic), direction of expression change (↑ upregulated, ↓ downregulated), sample source (e.g., plasma, serum, exosomes, skeletal muscle, or whole blood), and the original reference reporting the observation. This curated catalog provides a literature-based reference framework to contextualize exercise-associated miRNAs and to facilitate comparison with candidate miRNAs identified using the nf-Sarcopipe workflow.
